# Optimal Cancer Evasion in a Dynamic Immune Microenvironment

**DOI:** 10.1101/2022.08.03.502723

**Authors:** Jason T. George, Herbert Levine

## Abstract

The failure of cancer treatments, including immunotherapy, continues to be a major obstacle in preventing durable remission. This failure often results from tumor evolution, both genotypic and phenotypic, away from sensitive cell states. Here, we propose a mathematical framework for studying the dynamics of adaptive immune evasion that tracks the number of tumor-associated antigens available for immune targeting. We solve for the unique optimal cancer evasion strategy using stochastic dynamic programming and demonstrate that this policy results in increased cancer evasion rates when compared to a passive, fixed strategy. Our foundational model relates the likelihood and temporal dynamics of cancer evasion to features of the immune microenvironment, where tumor immunogenicity reflects a balance between cancer adaptation and host recognition. In contrast with a passive strategy, optimally adaptive evaders navigating varying selective environments result in substantially heterogeneous post-escape tumor antigenicity, giving rise to immunogenically hot and cold tumors.

## 1. Introduction

Cancer dynamics, encompassing both genotypic evolution and phenotypic progression, lies at the heart of treatment failure and disease recurrence, and therefore represents a significant and stubborn therapeutic hurdle. Prior research efforts have made substantial progress in detailing the mathematics of acquired drug resistance (Iwasa et al., 2006; Michor et al., 2004; Komarova, 2006) and the complementary roles of phenotypic and genotypic changes (Gupta et al., 2019). Recently, there has been much renewed interest in therapies that utilize the adaptive immune system to confer durable remission (Couzin-Frankel, 2013; Waldman et al., 2020). These latter breakthroughs have generated considerable interest in quantifying the cancer-immune interaction (Mayer et al., 2019; Sontag, 2017; George et al., 2017). As with targeted therapeutic resistance via compensatory evolution or adaptive rewiring (Bergholz and Zhao, 2021), tumors can similarly evade the immune system via either elimination or down-regulation of tumor-associated antigens (TAAs) normally detectable by the T cell repertoire (Rosenthal et al., 2019). However, several key features distinguish immune-specific evasion from classical drug resistance (Komarova, 2006). Dynamical changes in cancer genotypes and phenotypes, while problematic for conventional therapies, create additional TAAs that may subsequently be recognized by distinct T cells (Yarchoan et al., 2017). Thus, the evolving diversity of the T cell repertoire, consisting of billions of unique clones each with a distinct T cell receptor, provides adaptive immunity and immunotherapy the unique advantage of repeated tumor recognition opportunities (George and Levine, 2021; Lakatos et al., 2020; Qi et al., 2014), making long-term evasion more challenging.

Previous research efforts have investigated the diversity of evolutionary trajectories and the extent of cancer-immune co-evolution occurring in early disease progression (George and Levine, 2018, 2020). These works were based on increasing evidence of significant and sustained tumor evolution driven by immune surveillance (Turajlic et al., 2018; Jamal-Hanjani et al., 2017). Immunosurveillance via distinct T cell clones imposes an adaptive, stochastic recognition environment on developing cancer populations (Desponds et al., 2016) that co-exist with the immune system over large time scales (Turajlic et al., 2018), thereby motivating the need for a more complete understanding of the interplay between immune recognition and cancer evolution for effective therapeutic design. In addition to parsing this complexity, the precise extent to which a cancer population may *actively* evade repeated immune recognition attempts is at present unknown. Previous modeling efforts have assumed that cancer adaptation occurs passively, i.e. without behavior predicated on knowledge of the current immune microenvironment (IME). However, it is well-known that cancer populations commonly undergo phenotypic changes capable of altering their immunogenicity (Tripathi et al., 2016); these changes could be coupled to sensing of the IME in a manner similar to cancer mechanical, chemical, and stress sensing (Lee et al., 2019; Damaghi et al., 2013; Rosenberg, 2001). Moreover, direct experimental evidence demonstrates genetic adaptation in bacterial systems capable of sensing stress and consequently varying the per-cell mutation rate (Al Mamun et al., 2012; Rosenberg and Queitsch, 2014); there appear to be similar stress pathways in cancer (Bindra et al., 2007). Therefore, an alternative to passive evolution is for cancer populations to actively sense and evade recognition in the current environment en route to metastasis in a manner that maximally benefits survival, which we refer to henceforth as the ‘optimal escape hypothesis’. Understanding the extent and associated features of optimized tumor evasion is a crucial first step to identifying the best therapeutic approach, particularly for T cell immunotherapies that may be temporally varied.

Here, we introduce a mathematical framework, which we call ‘Tumor Evasion via adaptive Antigen Loss’ (TEAL), to quantify the aggressiveness of an evolutionary strategy executed by a cancer population faced with a varying recognition environment. This framework enables a dynamical analysis of both passive and optimized evasion strategies. The TEAL model describes a discrete-time stochastic process tracking the number of targets available to a recognizing adaptive immune system. We apply dynamic programming (Bellman and Dreyfus, 1959; Ross, 2014) in order to solve the corresponding time homogeneous Bellman equation detailing the tumor optimal evasion strategy for a specific example of the assumed penalty for attempting to avoid immune detection. In doing so, we obtain an exact analytical characterization of the evasion policy that maximizes long-run population survival, which we show is the unique solution. We can then quantify the enhancement in survival for optimal treats relative to their passive counterparts under a variety of temporally varying recognition environments. Surprisingly, we find that optimized strategies exhibit substantial diversity in their dynamical behavior, distinguishing them from threats with a fixed evolutionary strategy. Notably, immune recognition efficiency and the IME microenvironment are predicted to influence the likelihood for tumors to either accumulate or lose therapeutically actionable TAAs prior to their escape. The TEAL model represents a first attempt to explicitly represent – and in the future test – the optimal escape hypothesis, in order to frame cancer evasion as a dynamic and informed strategy aimed at maximizing population survival.

### Model Development

In greatest generality, our model consists of an evading clonal population that may be targeted over time by a recognizing system. We assume henceforth that the recognition-evasion pair consists of the T cell repertoire of the adaptive immune system and a cancer cell population, recognizable by a minimal collection of *s_n_* TAAs present on the surface of cancer cells in sufficient abundance for recognition to occur over some time interval *n*. Our focus is on a clonal population, recognizing that subclonal TAA distributions in this model may be studied by considering independent processes in parallel for each clone.

Experimental evidence and prior modeling suggests that tumors may be kept in an ‘equilibrium’ state of small population size prior to either escape or elimination, with repeated epochs of recognition and evasion (Dunn et al., 2004; Turajlic et al., 2018; George and Levine, 2020). We adopt a coarse-grained strategy and assume that during each each epoch, the immune system has an opportunity to independently recognize each of the *s_n_* TAAs with probability *q*, and also the cancer population can lose recognized TAAs, each with probability *π_n_*. This latter probability is either fixed or chosen by the cancer population using information available in the current period. If the immune system cannot detect any of the available TAAs in a given period, then the cancer population escapes detection. On the other hand, if *r_n_* > 0 antigens are detected by the adaptive immune system in this time frame, then the cancer population is effectively targeted. This leads to extinction of the cancer population unless it is able to lose each of the *r_n_* recognized antigens during the same period. This loss of recognition would presumably arise in a subpopulation which would then expand at the expense of the successfully targeted cells. If evasion balances recognition and all detected antigens are lost, then the process repeats in the next period with a new number of target antigens given by a state transition equation

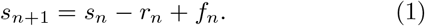

The last term *f_n_* represents the addition of new TAA targets for the next period. We shall refer to *f_n_* as the (inter-temporal) penalty term, the idea being that changes that lead to antigen loss will out of necessity give rise to the creation of new TAAs, in the form of either over-expressed/mislocalized self-peptides or tumorneo-antiegens.

The model therefore defines a discrete time process that involves changes to both the tumor and the immune system. The process ends in cancer elimination if the cancer population is unable to match all of the *r_n_* recognized antigens at any period. The process ends in cancer escape if at any period the number of recognized antigens is zero (*r_n_* = 0). This framework mirrors the outcomes resulting from known tumor-immune interactions, a process that leads via immunoediting to cancer escape, elimination, or equilibrium (Schreiber et al., 2002; Dunn et al., 2002, 2004; Koebel et al., 2007). Here, tumor antigenicity is represented by the total number of post-escape TAAs. We do not distinguish between different types of TAA loss, which may occur through a number of mechanisms including somatic mutation, epigenetic regulation, or phenotypic alteration.

#### 1.1. Passive evader

In the passive case, the cancer population does not change its evasion rate so that *π_n_* = *p* is fixed and independent of any of the parameters governing the recognition landscape. For this case, we shall also use the simple assumption that the penalty term *f* is a fixed constant.

#### 1.2. Optimal evader

In the optimized case, *π_n_* is chosen in order to maximize the overall evasion probability using date-*n* information. We assume that *s_n_* the number of TAAs as well as *r_n_* the size of the recognized subset is knowable by the cancer prior to strategy selection. In addition, we postulate that the inter-temporal penalty scales directly with *π_n_*, a reasonable assumption given, for example, the direct relationship between mutagenesis and passenger mutation accumulation (Pon and Marra, 2015; McFarland et al., 2014). While many functional forms of *f_n_* (*π_n_*, *r_n_*, *s_n_*) would be reasonable, we assume in general that the penalty is *π_n_*-affine:

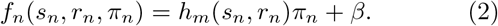

To make our system analytically solvable, we use a specific choice in which *h_m_* scales monotonically as a function of both *r_n_* and *s_n_* and *h_m_*∝*r_n_* in the large *r_n_* limit (see Sec. S3). We consider various assumptions for *β*. Since the number of recognizable (and thus actively targeted) TAAs reflect, all else being equal, an active immune microenvironment (IME) hostile to cancer, we assume that subsequent evasion penalties are dependent on the current level of immune detection, thereby taking into account the increased cost of surviving in, for example, an inflammatory IME. The temporal dynamics of the TEAL process are illustrated in Fig. 1A.

**Figure 1:**
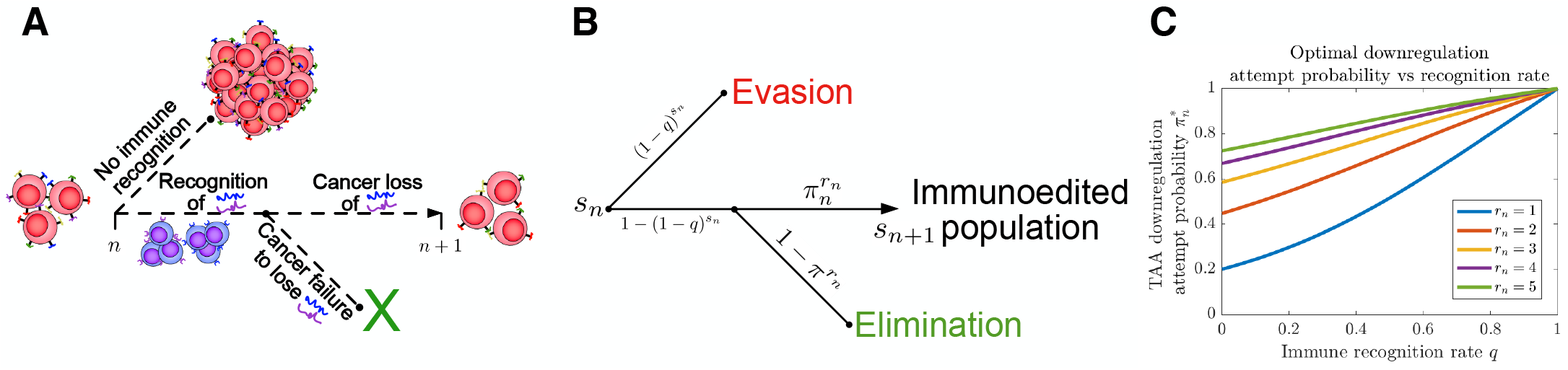
TEAL model. (A) Illustration of tumor antigen detection and down-regulation in the TEAL model of cancer-immune interaction; (B) The directed graph with nodes representing the states of the TEAL model and edges labeled based on the probability of their occurrence. Both evasion and elimination are absorbing states, and the tie state results in repeated interaction; (C) Plots of single-period cancer optimal evasion rates *π*\* given by Eq. 8 are plotted as a function of recognition rate *q* for various numbers of recognized antigens 0 < *r_n_* ≤ *s_n_* with *s_n_* = 5.

#### 1.3. Varying environments

Using the above framework, we subject both passive and active cancer evasion tactics to temporally varying recognition profiles. We partition pre-escape dynamics into four cases based on immune recognition *q* and evasion penalty *β*, from which we characterize the distribution of escape time, cumulative mutational burden, and predicted post-escape tumor immunogenicity.

## Results

The following section presents the main findings of our analysis (full mathematical details are provided in the SI). For *s_n_* available and *r_n_* recognized TAAs, we have that *r_n_* ~ Binom(*s_n_*, *q*). Conditional on recognition (*r_n_* > 0), the number of down-regulated antigens, *ℓ_n_*, is given by *ℓ_n_* ~ Binom(*r_n_*, *π_n_*). Recognition therefore occurs with probability 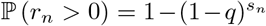. In a similar manner, non-elimination occurs following recognition with probability 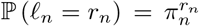. A decision tree for the TEAL process is illustrated in Fig. 1B.

### Passive evasion strategy

Here, *π_n_* = *p* is fixed. It can be shown (Sec. S2.4) that the dynamics of Eq. 1 in the passive case have exact mean evolution dynamics conditioned on non-escape/non-elimination as

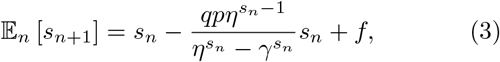

where *η* ≡ 1 – *q*(1 – *p*) is the probability of a tie for a single TAA given the existence of at least one available TAA. These dynamics may be approximated by

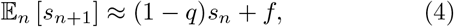

where 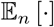 is the conditional expectation with respect to date-*n* information. The approximation given by Eq. 4 is a lower estimate of tumor antigenicity and is accurate as long as *p* and *q* are not both small and in particular for choices that give rise to large tie probability (Figs. S4-S7).

### Optimal evasion strategy

Here, *π_n_* is variable and the optimal choice will depend on *s_n_* and *r_n_*. The use of dynamic programming to address the optimal long-term evasion policy relies on a defined value function (Bellman and Dreyfus, 1959). We shall focus on the case where the cancer population is assigned normalized values of 1 at any period resulting in escape and 0 otherwise. The corresponding stationary Bellman equation takes the form

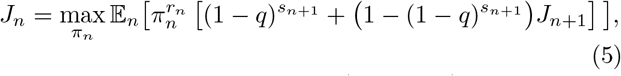

where the value function *J_n_*=*J*(*s_n_*, *r_n_*, *π_n_*) represents the maximal attainable value at period *n*; see SI Sec. S3 for full details. It can be shown that

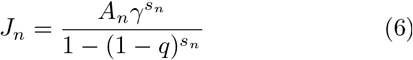

with

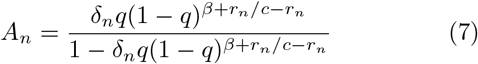

satisfies Eq. 5. Here, 0 < *δ_n_* ≤ 1 is a free parameter which varies inversely with the risk aversion of the evader (larger values imply a bolder strategy). One advantage of the dynamic programming approach is that it reduces an infinite-period optimization problem to a sequence of single-period optimizations. The corresponding optimal policy is given by the sequence

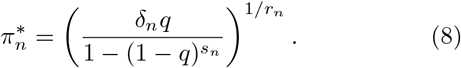

Plots of 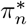 are given for various *r_n_* in Figs. 1C and S11. As expected, this closed-form strategy results in increased values for the attempt probability choice *π_n_*, which increase for increasing *q* and *r_n_*. We take *δ_n_* = 1 in subsequent analysis (so that the optimal strategy when *s_n_* = *r_n_* = 1 is 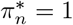).

### Active evasion strategies enhance population survival rates

For a fixed penalty, Eqs. 3 and 4 describe a mean-reverting process. Consequently, the mean number of TAAs approaches a stable equilibrium

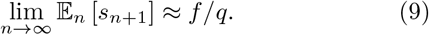

as long as the cancer neither escapes nor is eliminated. In the optimal case, a similar equilibrium value *s*_∞_ may be calculated:

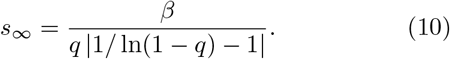

In this case, stability is more complex: If immune recognition is sufficiently effective, meaning *q* > *q*\* = 1 – *e*^−1^, then Eq. 10 is a stable equilibrium exhibiting mean reversion similar to that of the passive case. On the other hand, recognition impairment (*q* < *q*\*) gives rise to an instability, which results in a system harboring an initial number of targets *s*_0_ being driven either to escape if *s*_0_ < *s*_∞_ or to large accumulations (and likely elimination) if *s*_0_ > *s*_∞_ (Fig. S13).

We proceed by contrasting active and passive escape rates assuming no recognition impairment, and discuss the implications of immune impairment in a later section. Simulations of passive and optimized strategies with passive evasion rates matching mean optimal evasion rates 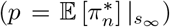 are compared in Fig. 2. Despite identical mean TAA evolution (Fig. 2A) and comparable intertemporal penalties, the optimized strategy results in substantially higher cancer escape probability (150%) compared to the passive case. Moreover, optimized strategies generate wider escape time distributions, thus illustrating an adaptive evader’s sustained effort to thwart elimination prior to escape (Fig. 2B).

**Figure 2:**
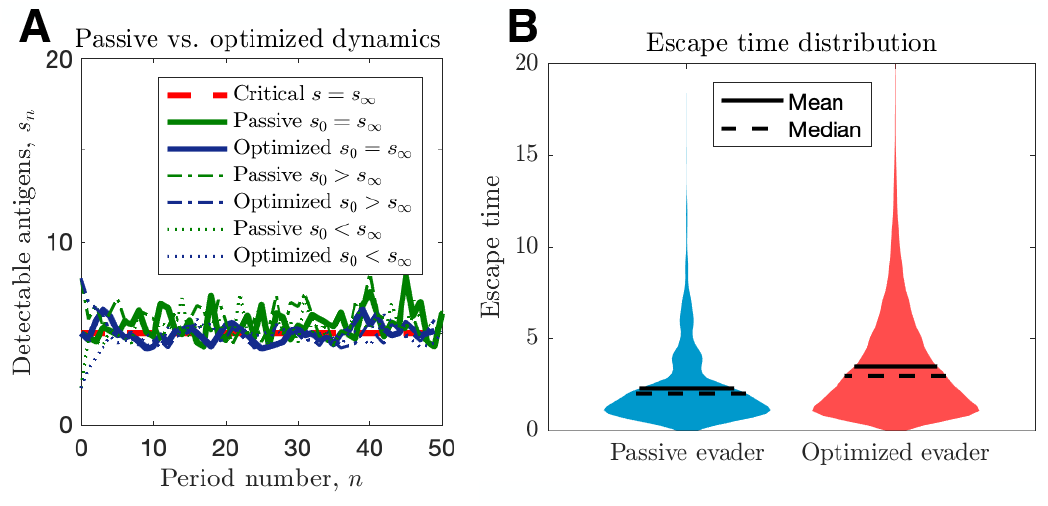
Passive and optimized evasion strategies against stationary threats. (A) Comparisons of the temporal dynamics of passive (green) and active (blue) strategies with parameter selections giving equal mean behavior. In the active case, *q*=*q*\*+0.1 yields stable dynamics, giving mean adaptive penalty *f*=3.44. In the passive case, *p*=0.90 was selected to match the mean optimal evasion rate and the expected *s_n_* of the active case. Also, *f*=4.39 and *β*=0.88 both chosen so that *s*_∞_=5, and the results plotted for *s*_0_∈{2, 5, 8}); (B) 10^6^ replicates of this process were used to calculate distributions of stopping times conditioned on escape. This distribution generates passive (resp. optimized) *p*_escape_ of 5.37·10^−3^ (resp. 8.44·10^−3^).

### Arbitrary recognition landscape

The above describes the dynamics of passive and optimized cancer co-evolution during adaptive immune recognition with constant governing parameters. We can more generally apply this approach to understand how an evasion strategy affects the likelihood and timing of cancer escape under a variety of temporally varying recognition landscapes. Such landscapes could, for example, be imposed by a clinician temporally modulating an immunotherpauetic intervention, and are routinely proposed in the setting of traditional therapies, where attempted strategies have included a variety of cyclical burst approaches (Foo and Michor, 2009; Eigl et al., 2005). A similar approach could be taken with regard to timing and dosage of adoptive T cell immunotherapy. An advantage of our dynamic programming approach is the ability to study optimal evasion strategies for arbitrary recognition landscapes (Fig. 3A). We simulate TEAL dynamics and find that optimized immune evaders are more successful in evading detection than their passive counterparts across various recognition landscapes (Fig. 3B). Evasion, when it occurs in the optimized case, does so largely after a sustained interaction with the recognizing threat (Fig. 3C). Collectively, our results detail the dynamics of sustained cancer-immune co-evolution via TAA loss in threats capable of adopting adaptive evasion strategies in the presence of complex treatment modulation (George and Levine, 2020; Turajlic et al., 2018).

**Figure 3:**
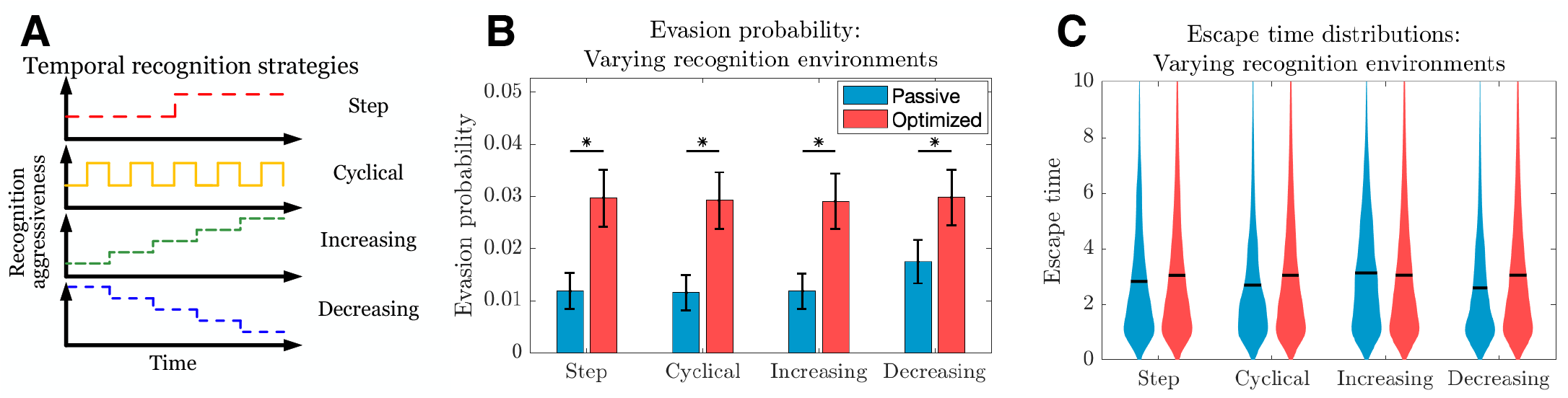
Passive and optimized evasion strategies for temporally varying recognition profiles. (A) Temporally varying recognition functions are selected and applied to threats employing passive (blue) and optimized (red) evasion strategies. (B) The mean and standard deviation of escape probabilities is compared across recognition profiles for each strategy (Pairwise significance was assessed using 2-sample t-test at significance *α*=0.05 with *p*<10^−5^). (C) Escape time distributions are generated for step, cyclical, increasing, and decreasing recognition environments (solid line: mean). In each case, mean inter-temporal penalties for passive (resp. optimized) evasion were 4.39 (resp. 4.75), and 10^3^ simulations of 10^3^ replicates each were used for statistical comparison; all samples were aggregated for escape time violin plots (solid line denotes mean).

### Optimal evaders under effective immune recognition accrue mutations at a fixed rate

One consequence of mean reversion is that the rate of mutation accumulation over time, λ(*n*), is linear in n (Sec. S3.4):

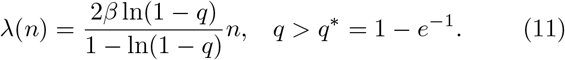

The prediction of constant accumulation is consistent with empirically observed cancer mutation behavior (Lawrence et al., 2013; Alexandrov et al., 2013). This is not what holds in the impaired case (as will be discussed later), thus suggesting that early cancer progression often proceeds in an environment with effective immune recognition. Additionally, our formula shows that larger mutation rates can be caused by large evasion penalties or by reduced immune recognition. Of course, the TEAL model does not consider any specific features that determine the values of the effective parameters. Instead, its utility is in quantifying the overall effect of reducing antigen detection resulting from, for example, transitions to an immune impaired microenvironment.

### Post-escape tumor antigenicity determined by a balance between recognition aggressiveness and local penalties in the immune microenvironment

The prior section related recognition and penalty to observed mutation rates. We now consider their combined effects on tumor immunogenicity following immune escape. The TEAL model represents immunogenicity by the number of available TAAs at the time of cancer detection, an important predictor of immunotherapeutic efficacy (Martin et al., 2016; Samstein et al., 2019; Goodman et al., 2017). We apply the TEAL model to simulate evading cancer populations, focusing exclusively on trajectories that result in tumor escape, to characterize the distribution of available TAAs. This is performed first for increasing immune recognition rates *q* (Fig. 4A) and then for increasing penalty term *β* (Fig. 4B). Our results demonstrate that larger penalties result in higher post-escape TAA levels, while efficient immune recognition depletes available TAAs. The presumptive reason for this latter observation is that escape in the presence of strong immune recognition biases the tumor to have low numbers of TAAs. This prediction agrees with recent empirical observations that strong immune selective pressure in early cancer development results in tumor neoantigen depletion and is prognostic of poor clinical outcome (Rosenthal et al., 2019; Lakatos et al., 2020).

**Figure 4:**
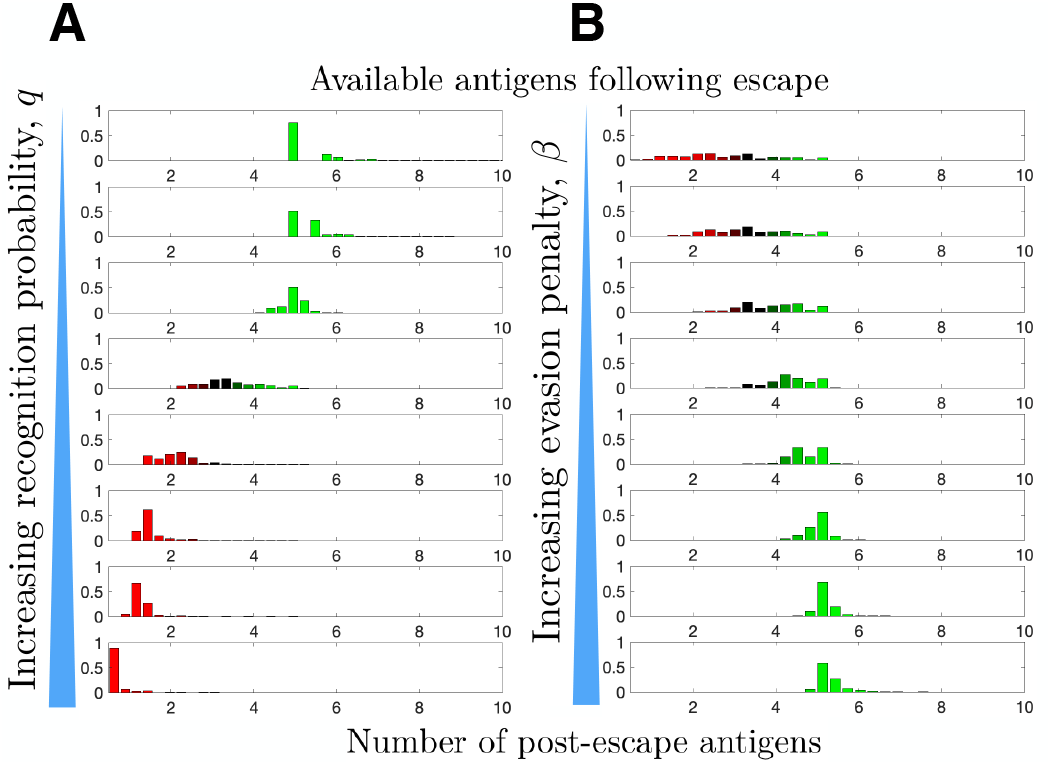
Distribution of available post-escape tumor antigens. The distribution of TAAs were estimated from simulations of optimized cancer evasion resulting in escape and plotted for (A) increasing recognition probability *q* ∈ {0.6, 0.65, 0.7, 0.75, 0.8, 0.85, 0.9, 0.95} and; (B) increasing evasion penalty *β* ∈ {0.10.20.30.40.50.60.70.8} (For (A), *β* = 0.59. For (B) *q* = 0.7 > *q*\*. In both cases, *s*_∞_ = 5 and *n* = 10^6^ simulations were performed for each histogram.

### Variation in the tumor microenvironment drives the generation of immune hot vs cold tumors under optimal evasion

In the passive evader case, antigenicity fluctuates around a stable equilibrium that varies directly with penalty and inversely with recognition (Sec. S2). The adaptive case (Sec. S3) gives rise to more complex behavior resulting from impairments in immune recognition or changes in penalty (Figs. S12, S13). These changes are important manifestations of disease progression, which may alter the immunogenic landscape via impairments in immune recognition, such as MHC downregulation, costimulation alteration, T cell exclusion, or the establishment of a pro-tumor IME, via for example M2 macrophage polarization (Liu et al., 2021; Goswami et al., 2017). Although many factors may affect recognition rates, for simplicity we shall refer to larger vs. smaller immune recognition rates q as *infiltrated* vs. *excluded*.

On the other hand, the generation of new TAA targets is expected to vary substantially across tumor type, for example due to differing somatic mutation rates. Within a given tumor subtype, variations in the hostility of the IME, resulting from a large variety of possible mechanisms (metabolic, mechanical, cytokine, environment) requires cancer populations to undergo greater degrees of adaptation to survive; in our approach, this greater degree of adaptation comes with a greater penalty. Consequently, we relate large vs. small local penalty terms *β* to *antitumor* vs. *pro-tumor* IMEs. Conceptually, the baseline state (infiltrated anti-tumor IME) may give rise to three alternative states (excluded anti-tumor IME, infiltrated pro-tumor IME, or excluded pro-tumor IME), based on progression.

Toward this end, we simulate the TEAL model under the above conditions and record post-escape TAA distributions. As already explained, our results predict that infiltrated (*q* > *q*\*) environments lead to an absorbing equilibrium state in the intervening period prior to escape, while exclusion (*q* < *q*\*) results in unstable equilibria. Interestingly, the sign of this equilibrium, and hence the longterm immunogenic trajectory, depends on the sign of *β* (Sec. S4.4). The baseline infiltrated anti-tumor case (*q* > *q*\*, *β* > 0) yields a positive and stable, mean-reverting TAA steady state, generating immunogenically ‘warm’ tumors. Excluded anti-tumor IMEs (*q* < *q*\*, *β* > 0) exhibit low recognition and large TAAs arrival, resulting in a unstable TAA steady state that leads to increased immunogenicity over time, resulting in ‘hot’ tumors. Furthermore, the infiltrated pro-tumor (*q* > *q*\*, *β* < 0) case demonstrates preserved recognition with low TAAs arrival and generates an unphysiological negative stable steady state, thereby predicting that trajectories reduce immunogenicity to zero over time, yielding ‘cold’ tumors. Lastly, excluded pro-tumor IMEs (*q* < *q*\*, *β* < 0), having compromises in both recognition and TAA arrival rate, result in an unstable state, above which trajectories accumulate additional TAAs over time, becoming immunogenically ‘hot’, and below which the populations are predicted to reduce the number of recognizable TAAs over time, becoming ‘cold’ (Fig. 5A,B). Substantial heterogeneity in the distributions of escape time predict sustained interactions in the unimpaired case (Fig. S14). Tumor exclusion leads to hot tumors so that escape, should it occur, must do so on average prior to the accumulation of many TAAs. Conversely, pro-tumor IME with immune recognition drives TAA depletion, so escape occurs relatively early. These results are summarized in Fig. 5C.

**Figure 5:**
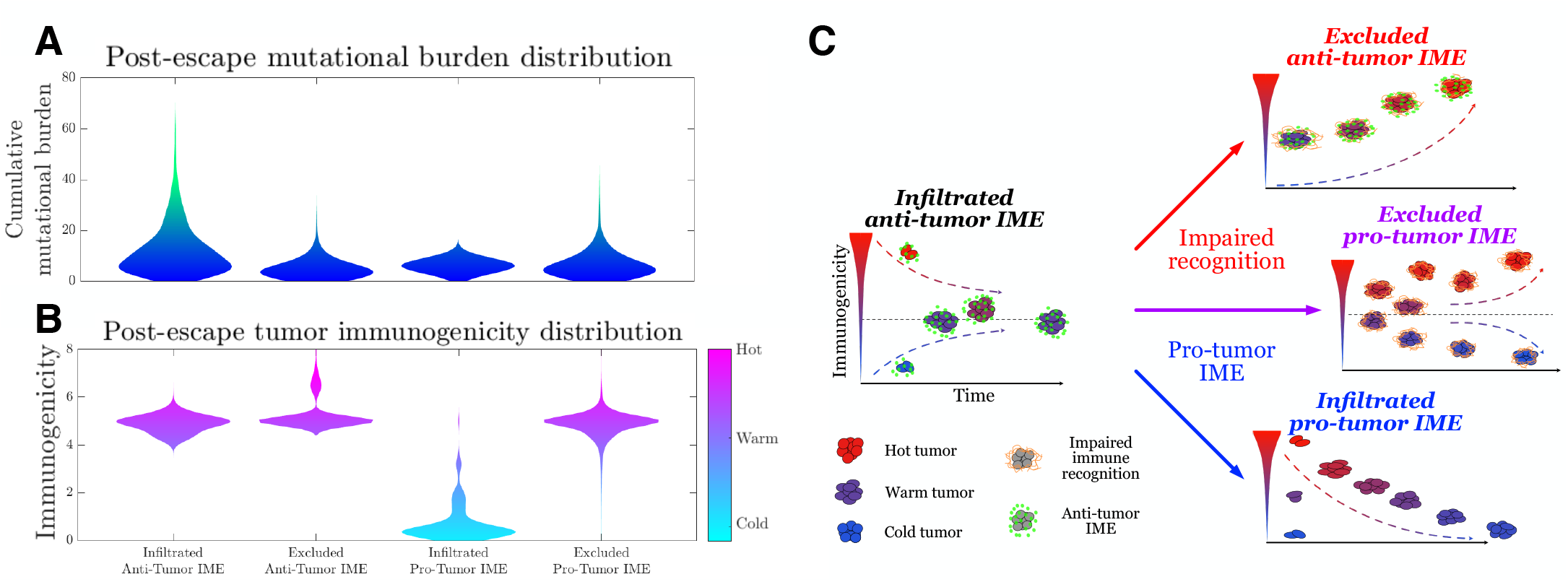
Active Evader dynamics. Violin plots of of the distribution of post-immune escape (A) cumulative mutation burden (B) post immunogenicity (available TAAs) assuming as a function of time for a variety of tumor IME conditions (Anti-tumor infiltrated: *q* = *q*\* +0.1, *β* = 0.529; Anti-tumor excluded: *q* = *q*\* – 0.1, *β* = 0.505; Pro-tumor infiltrated: *q* = *q*\* +0.1, *β* = −0.529; Pro-tumor excluded: *q* = *q*\* – 0.1, *β* = −0.505. In all cases, *β* chosen to give |*s*_∞_| = 3 (*s*_∞_ =−3 for the pro-tumor infiltrated case) giving strictly positive penalties. Simulations were run until *n* = 10^6^ escape events occurred for each case). (C) The number of recognizable TAAs over time along with equilibrium states are depicted assuming (left) anti-tumor IME, *β* > 0 and efficient immune recognition. Compromises in (top right) recognition, *q* < *q*\*; (bottom right) the establishment of a pro-tumor IME, *β* < 0, or; (middle right) both affect the predicted dynamical behavior of tumor immunogenicity.

## Discussion

The underlying evolutionary dynamics of adaptive populations lies at the heart of many important clinical challenges, including antibiotic resistance, acquired drug resistance, immunotherapy failure, and tumor immune escape. Quantitative analytic modeling will continue to provide improved insight into these complex issues by generating fast and affordable predictions and a convenient theoretical framework for hypothesis testing. To-date, virtually all of the current models of cancer evolution and the tumor-immune interaction have assumed passive acquired evolution without allowing the tumor to sense and optimally respond to the current fitness landscape in order to maximize future survival. The ‘optimal escape hypothesis’ is, in our opinion, worth exploring in light of the myriad examples of treatment failure and adaptive resistance.

Toward this end, we propose and analyze the TEAL model for studying and comparing passive and optimal escape mechanisms in the tumor-immune interaction. We focused our dynamic programming approach on a particular set of relations to provide analytical insight into this process. We do note, however, that the Bellman function approach to dynamic programming can be numerically implemented to obtain solutions for arbitrary functional forms of the penalty function, thereby enabling analysis of more complex assumptions where analytic progress becomes intractable. As expected, threats adopting optimal evasion strategies largely out-perform their passive counterparts by increasing the rate of immune escape over prolonged cycles of cancer-immune co-evolution. In the setting of the tumor-immune interaction, the resulting TAAs available for targeting, a proxy for clinical post-detection immunotherapeutic efficacy, is augmented when cancer populations accrue large penalties for evasion and, perhaps surprisingly, when immune recognition is impaired.

Evasion dynamics of passive and active evaders are similar in some ways while different in others. Similarities include the mean-reverting stationary dynamics of both strategies under efficient immune recognition. However the TEAL model predicts, for adaptive threats in an excluded pro-tumor IME, the emergence of an unstable state resulting in either accrual or depletion of TAAs in a manner that depends on the current TAA abundance. This splitting behavior into ‘hot’ and ‘cold’ tumors offers insight into the microenvironmental features generating spatial immunogenic diversity within solid tumors and is consistent with prior observations (Huss et al., 2021; Jia et al., 2022; Meiller et al., 2021; Lakatos et al., 2020). This argues that TAA-depleted tumors share in common the tendency for their evasion strategies to incur less antigenic penalties. Our results suggest the possibility of altering the tumor IME to increase the immunogenicity of immune-cold tumors by making evasion more costly, in a manner reminiscent of mutational meltdown (Gabriel et al., 1993). We remark that these dynamics are worth considering in the case of adoptive T cell-based immunotherapies, marked by their potential for exerting substantial co-evolutionary pressure on a developing malignancy (George and Levine, 2021). We also predict that impaired immune recognition leads to TAA accumulation, consistent with experimental observations in lung cancer wherein patients with HLA loss of heterozygosity harbored larger mutational burdens, an indirect measure of TAAs of our model (McGranahan and Swanton, 2017). Lastly, active evader variable mutation rates also distinguish this case from passive evaders with fixed mutation rates, and this feature is analogous to that observed in bacterial colonies faced with antibiotic selective pressure (Windels et al., 2019).

More generally, the TEAL framework provides a mechanistic basis for several empirical observations. First, our results would suggest that the lower observed TAA availability of hematological malignancies vs. immune-protected solid tumors, such as melanoma (Lawrence et al., 2013), occur as a result of greater immune accessibility and possible immunoediting of liquid cancers. Second, our model predicts enhanced immune interactions, both natural and treatment-derived, resulting from increasing the cost of immune evasion in the evading cancer population in order to enrich the TAAs following escape. This supports the utility of neo-adjuvant radiation therapy (McGranahan et al., 2016) or chemotherapy (Mouw et al., 2017) in inducing immunogenicity. Orthogonal efforts to quantify cancer evolution have similarly predicted the benefit of larger evasion rates resulting in mutational meltdown (McFarland et al., 2014). Integrated together, the TEAL model can predict the balance of generated TAAs given the relative influences of recognition and evasion penalty.

Tumor antigen depletion is a concerning consequence of immunotherapy, since increased recognition is desirable and required for tumor elimination. In solid tumors, one contributor to this problem is T cell exclusion. (Pai et al., 2020). However, should effective treatment and robust tumor recognition lead to relapse, the resulting tumor has a greater chance of being TAA-depleted (Rosenthal et al., 2019). Other strategies that fall in this group include those that effectively reduce recognition, like the presence of T-regulatory cells. Our results suggest that this detrimental effect of targeting can be offset by increasing the ‘hostility of the IME’. Strategies encouraging making tumor adaptation more penalizing, such as fostering an anti-tumor environment by, for example, M1 macrophage polarization, or the inactivation of Tumor associated macrophages (Liu et al., 2021; Goswami et al., 2017).

Of course, this foundational model is not without limitation. At present, we have assumed that the recognition agent is not employing an optimized strategy informed by optimal cancer evasion. Instead, we have detailed our results for arbitrarily imputed recognition landscapes, which is useful for predicting the response of an aggressive evader like cancer to particular immunotherapeutic interventions, such as hematopoietic stem cell transplant and adoptive T cell therapy, where the clinician has temporal control over treatment. Identification of such optimal treatment strategies upon quantification of disease evasion aggressiveness, is of paramount importance. Furthermore, we at present consider a cancer population with core set of clonal TAAs. Distinct tumor subclones, each with a different pattern of TAA availability, would necessitate a series of TEAL models tracking their independent evolution in parallel. Lastly, cancers characterized by co-evolutionary dynamics resulting in large variability in population size prior to escape or elimination would require in general that recognition and evasion parameters depend on the current period. While possible to incorporate, we have for foundational understanding assumed these to be constant.

We detailed strategies that affect the number of TAAs present following escape. In addition to quantity, variations in individual TAA antigenicity could affect overall immunogenicity, but we do not as yet take this into account. In future work, individual antigenicities could be built in by allowing individual TAA contributions to *s_n_* and *q* to depend on the particular TAA. Many additional features contribute to the immune landscape. Here, we focused on TAA availability and effects of general immune recognition rates and IME hostility on TAA accural. Future efforts may incorporate additional cancer-specific features, including antigen presentation, immunomodulatory gene expression, and measured immune signatures present in the IME.

These optimized dynamics are proposed in absence of the precise mechanistic details of cancer decision-making. Further studies linking changes in the evasion rates to cell signaling are necessary next steps at elucidating a possible mechanism of optimal evasion. Our framework serves as a tool for evaluating the extent of evasion aggressiveness in a variety of observed disease contexts, including cancer. Differentiating dynamics of passive and adaptive evasion mechanisms is a first step to understanding this difference, its importance underscored by the large implications such an understanding would have on our approach to treatment.

The TEAL model represents a framework broadly applicable for studying population behavior consistent with optimized collective decision-making, and subsequent experimental validation or refutation is of highest priority. Future direction aims to apply this framework for personalizing optimal interventions which maximize disease elimination probabilities. Consequently, stochastic analysis and optimal control theory are indispensable tools for better understanding the complex cancer-immune interaction. Defeating an evolving cancer population has provided a persistent challenge to researchers and clinicians, with the majority of progress heralded by fundamental discoveries on cancer behavior, and additional insights require a more detailed understanding of cancer evasion. The possibility that cancer population-level strategies are somewhat informed to the present recognition threat would have a radical effect on our own optimal treatment approach.

## Methods

A complete description of all mathematical details may be found in the SI Appendix.

## Author’s Contribution

J.T.G. conceived of and designed the research, analyzed the data, contributed new analytic equations, and wrote the manuscript. H.L. designed research, analyzed the data, and wrote the manuscript. Both authors have approved the final article.

## Acknowledgments

J.T.G. would like to thank Kerry E. Back, Philip A. Ernst, Thomas J. George, and Richard A. Tapia for their helpful discussions on stochastic dynamic programming and optimization. J.T.G. was supported by the Cancer Prevention Research Institute of Texas (RR210080). J.T.G. is a CPRIT Scholar in Cancer Research. H.L. is supported by the National Science Foundation (NSF) grant NSF PHY-2019745.

## Appendix

Supplementary information with full mathematical details are provided in the attached document.

