## Supplementary Information for "Optimal Cancer Evasion in a Dynamic Immune Microenvironment"

August 3, 2022

### S1 Overview

Here, we present the full mathematical details of the Tumor Evasion via adaptive Antigen Loss (TEAL) model, characterizing the stochastic evasion of an evolving population to an varying recognition environment. The Sec. S2 considers dynamics under an adaptive treatment strategy like T-cell immunotherapy therapy for a passive evader, and Sec. S3 considers the same for a threat which may optimize its evasion strategy. In each case we predict the effects of various interventions and enhancements of each therapeutic strategy. This is, to our knowledge, the first attempt to model a threat like cancer taking an informed, active evasion and evolutionary strategy that optimizes the chance of future immune evasion, utilizing a policy similar to, for example, a rational decision-maker in a market.

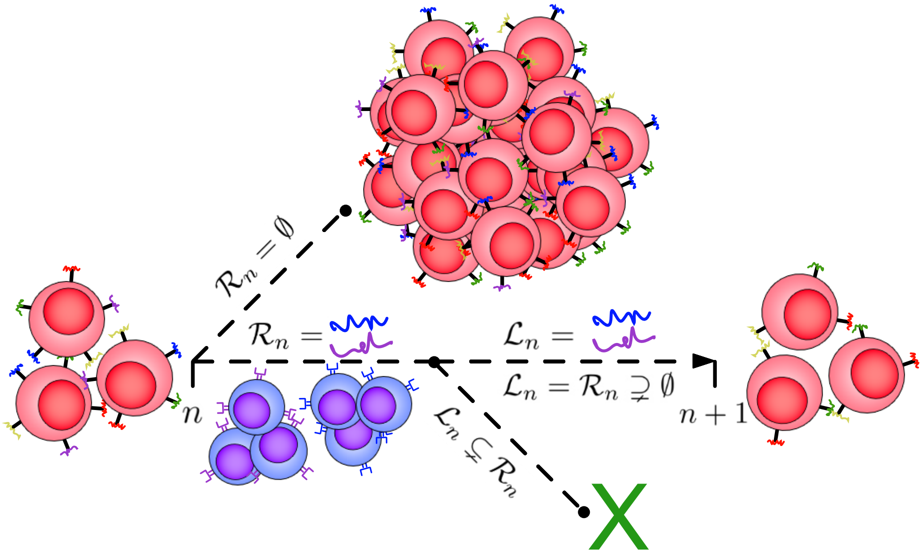


Figure S1: Illustration of the TEAL model. A population of cancer cells (red) possesses an initial minimal collection of surface antigens which may be identified and targeted by the immune system (colored antigens). If the immune system is unable to recognize any of these antigens ($\mathcal{R}_n = \emptyset$), then the cancer grows unchecked to escape size and the process ends. Alternatively, if the immune system (blue cells) recognizes a subset of detectable antigens ($\mathcal{R}_n \neq \emptyset$), then the cancer population must evolve by generating a clone which has avoided detection through the targeted antigen ($\mathcal{L}_n = \mathcal{R}_n$). If this occurs, then the cancer population survives to the next period, otherwise the population becomes extinct.

### S2 Passive Evader in an Adaptive Environment

Let $\mathcal{S}_n$ denote the set of tumor antigens recognizable by the immune system and present at period $n$ on a population of cancer cells, and let $s_n = |\mathcal{S}_n|$ count their number ($|\mathcal{A}|$ denotes the cardinality of set $\mathcal{A}$). From one period to the next, each of the $s_n$ detectable antigens may be independently and identically detected by the immune system with probability $q$ per antigen. We let $\mathcal{R}_n \subseteq \mathcal{S}_n$ denote the collection of antigens that are recognized by the immune system at time $n$. As the immune system targets and begins to eliminate cells via the $\mathcal{R}_n$ antigens, the cancer population has an opportunity to lose or down-regulate each of the $r_n = |\mathcal{R}_n|$ recognized antigens with a similar independent and identical manner, having probability $p$ per antigen. We denote the collection of antigens that are lost by the cancer population at time $n$ by $\mathcal{L}_n \subseteq \mathcal{S}_n$. We track the number of recognized and lost antigens at time $n$ by $r_n$ and $\ell_n = |\mathcal{L}_n|$, respectively, so that $\ell_n \leq r_n \leq s_n$.

The system evolves as follows: If $\mathcal{R}_n = \emptyset$, then the immune system is unable to recognize any tumor antigen at time $n$ and so the process ends in cancer escape. Since in this case the immune system *loses*, we denote this event by $L_n$. If $\mathcal{R}_n \neq \emptyset$, then the immune system recognizes the threat by at least one TAA and one of two outcomes results: The first possibility is that the cancer population successfully down-regulates or loses all of the targeted antigens, expressed as $\mathcal{L}_n = \mathcal{R}_n$, and survives to the next time step. We call this a tie and denote the event by $E_n$. Alternatively, the cancer population is unable to lose every recognized antigen and subsequently becomes eliminated. This means the immune system has *won* so we denote this event by $W_n$. Although the recognition and evasion probabilities may in general be clonally and temporally dependent, we assume fixed probabilities for the recognition, $q$, and evasion, $p$, of individual antigens. In the event of a tie, $s_n - r_n$ antigens remain, and a possibly noisy penalty term $f_n$ is added to reflect random production of new antigens as the population evolves. For simplicity, we assume the $f_n$ are a sequence of independent, identically distributed (IID) random variables with mean $f$. While it is in general possible that the distributions of $r_n$ and $\ell_n$ be both state- and time-dependent, we focus on the foundational example above.

This process is identical to the following game between two players, hereafter referred to as the 'Recognizer' (immune system) and the 'Evader' (threat): The Recognizer starts off with a collection, $\mathcal{S}_0$, of $s_0$ coins and begins her turn by flipping each coin with IID success probability $q$. If she has no success ($\mathcal{R}_0 = 0$), she loses (denoted by event $L_0$) and the game ends. If $r_0 > 0$ of her coins land on heads, then the next turn goes to the Evader, who proceeds to flip his $r_0$ coins with IID success probability $p$ in an attempt to match the Recognizer's successful coin flips. The Evader must succeed in all coin flips ($\mathcal{L}_0 = \mathcal{R}_0$) for the turn to end in a tie (event $E_0$). Otherwise, he loses and the game ends with a Recognizer win, (event $W_0$). If a tie occurs then both players restart the game, but only after the removal from $\mathcal{S}_0$ of the $r_0$ coins that landed on heads for both players as well as the addition of a random number $f_0$ of new coins. The Evader wins by default if a new turn begins and there are no longer any remaining coins to flip.

#### S2.1 Tie probability

It is immediately apparent that this game is unfair to the Evader if $s_0$ is much larger than 1, unless the recognition probability $q$ is low and the evasion probability $p$ is high. We motivate the following analysis with this in mind, and proceed to characterize the dynamics of this stochastic process. Clearly, the number of recognized and lost antigens during each period is binomially distributed, their respective distributions given by

$$r_n \sim \text{Binom}(s_n, q); \qquad \ell_n \sim \text{Binom}(r_n, p). \tag{S1}$$

The event of a tie (non-escape and non-extinction) may be written as

$$E_n = [\mathcal{L}_n = \mathcal{R}_n \supsetneq \emptyset] = [\ell_n = r_n > 0]. \tag{S2}$$

One might expect that the number of antigens lost at time $n$ is affected by knowledge of whether or not the game continues to be played. The distribution of $\ell_n$ conditioned on a tie may be characterized by

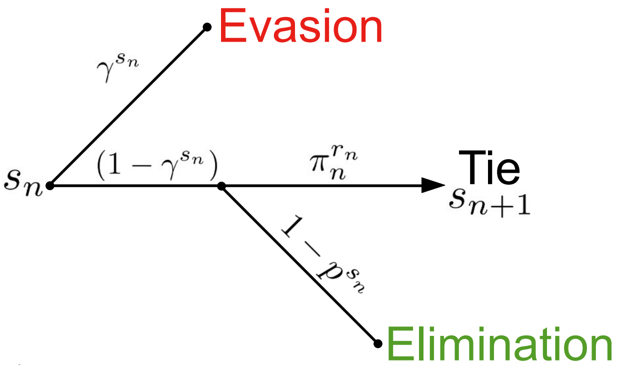


Figure S2: Passive Evader outcome tree.

conditioning on the number of recognized antigens at time $n$. To this end, let $F_{n,r} = [r_n = r]$ denote the event that $r$ antigens are recognized at period $n$, with

$$\mathbb{P}(F_{n,r}) = \binom{s_n}{r} q^r (1-q)^{s_n - r}. \tag{S3}$$

We remark that events $\{F_{n,r}\}_r$ are disjoint and exhaustive; in other words, for sample space $\Omega$,

$$\bigcup_{r=0}^{s_n} F_{n,r} = \Omega, \qquad F_{n,i} \cap F_{n,j} = \emptyset, \text{ for } i \neq j. \tag{S4}$$

Additionally, we note that a tie cannot occur if no antigens are recognized (i.e. $F_{n,0} = [\mathcal{R}_n = \emptyset]$). Lastly,

$$\mathbb{P}(E_n \mid F_{n,r}) = p^r, \tag{S5}$$

since if $r$ antigens are recognized then $\mathcal{L}_n = \mathcal{R}_n$ occurs if and only if each of the $l_n = r_n$ recognition positions are exactly matched with $r_n$ evasions. We will make use of the following variables to simplify subsequent results:

$$\eta \equiv (1-q) + qp = \big[1 - q(1-p)\big] \tag{S6}$$

and

$$\gamma \equiv 1 - q. \tag{S7}$$

Here, $\eta$ may be interpreted as the probability of the complement of the following event: "recognition occurs without matched evasion for a single antigen". In other words, $\eta$ is the probability of a tie at a single antigen position given the existence of at least one antigen. This event occurs in one of two disjoint ways for a single antigen: either there is no recognition, and so a tie occurs regardless of evasion, or there is recognition that must also be matched by evasion. The joint distribution of recognized and lost antigens is given by probability mass function

$$\begin{aligned} m(r,l) &= \mathbb{P}([r_n = r] \cap [\ell_n = \ell]) \\ &= \mathbb{P}(\ell_n = \ell \mid r_n = r)\ \mathbb{P}(r_n = r) \\ &= \binom{r}{\ell} p^\ell (1-p)^{r-\ell} \cdot \binom{s_n}{r} q^r (1-q)^{s_n - r}. \end{aligned} \tag{S8}$$

The probability that a tie occurs and the game continues at period $n$ is given by:

$$\begin{aligned}\mathbb{P}(E_n) &= \sum_{r=1}^{s_n} m(r,r) \\ &= \sum_{r=1}^{s_n} \binom{s_n}{r} (pq)^r (1-q)^{s_n - r} \\ &= (1-q)^{s_n} \left[ \left( \frac{q - pq - 1}{q-1} \right)^{s_n} - 1 \right] \\ &= \left[1 - q(1-p)\right]^{s_n} - (1-q)^{s_n} \\ &= \eta^{s_n} - \gamma^{s_n}, \end{aligned} \tag{S9}$$

which is equal to the probability of a tie occurring at every position minus the probability that all of the $s_n$ antigens are not recognized, since at least one recognized antigen is required for a tie to occur.

### S2.2 Break-even probability

The process is usually more favorable for the Recognizer. The Recognizer loses at period $n$ if there are zero recognition events, and this occurs with probability

$$\mathbb{P}(L_n) = \gamma^{s_n}. \tag{S10}$$

The Recognizer wins at period $n$ if she does not lose or tie, which occurs with probability

$$\mathbb{P}(W_n) = 1 - \Big( \mathbb{P}(E_n) + \mathbb{P}(L_n) \Big) = 1 - \eta^{s_n}. \tag{S11}$$

If $q$ and $s_n$ are given, then the evasion probability $p$ required for equal probabilities of Recognizer failure and success, or the *break-even probability*, is given by

$$p_{\text{even}} = \frac{(1-\gamma^s)^{1/s} - \gamma}{1-\gamma}, \tag{S12}$$

and exists whenever $p_{\text{even}} > 0$. We plot $p_{\text{even}}$ as a function of recognition probability $q$ for various numbers of TAAs, $s$ (Fig. S3A). The "fair-game" line indicates where the break-even evasion probability is always equal to the recognition probability. Regions where the break-even probability localizes above the fair-game line favor the Recognizer since there the evasion rates $p$ must be higher than recognition rates $q$ for the game to be fair. Alternatively, areas below the break-even curve favor the Evader. It is clear from Fig. S3B that the process favors recognition for a majority of parameter choices $(p, q)$ in all cases except for when $s = 1$. Thus, the process is largely unfair and mostly favors the Recognizer over the Evader when $p = q$ so long as $s$ is not small. In order for the Evader to have a reasonable chance of success, either the evasion probability must be very large, or the number of TAAs must remain small.

### S2.3 Distribution of lost antigens

The process transitions at period $n$ if and only if a tie occurs, which means that the number of lost antigens match those recognized and are strictly positive. In other words,

$$E_n = [\ell_n = r_n > 0]. \tag{S13}$$

The survival probability as a function of $q$ and $p$ are plotted for various choices of $s$ in Fig. S4. From this, we find that ties occur with high probability for large evasion rates, $p$, as well as for recognition rates $q$ that vary inversely with the number of recognizable antigens. This coincides with conditions that do not disadvantage the Evader so that the tie probability is maintained. We remark that recognition and evasion rates in general vary with the immune microenvironment (IME). We shall subsequently restrict our attention to large recognition probabilities ($p > 1/2$).

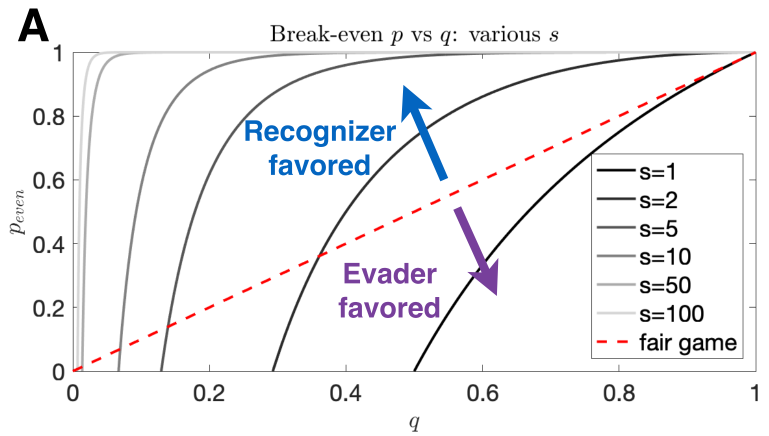


B

| s | Minimal break-even q | Area under break-even curve |
|---|---|---|
| 1 | 0.500 | 0.307 |
| 2 | 0.293 | 0.557 |
| 3 | 0.206 | 0.678 |
| … | … | … |
| 10 | 0.067 | 0.891 |
| … | … | … |
| 100 | 0.007 | 0.989 |

Figure S3: Break-even evasion probability as a function of recognition probability. (A) The break-even probability is given as a function of recognition probability for various numbers of recognizable antigen. The red dashed line illustrates a fair-game, where an equal proportion of parameters favors the Evader and Recognizer. Recognition (resp. evasion) is favored in systems for which the area below the $p_{\text{even}}$ curve is large (resp. small); (B) values for the minimal evasion probabilities allowable for the existence of a break-even probability as well as area under the break-even curve for various antigen numbers.

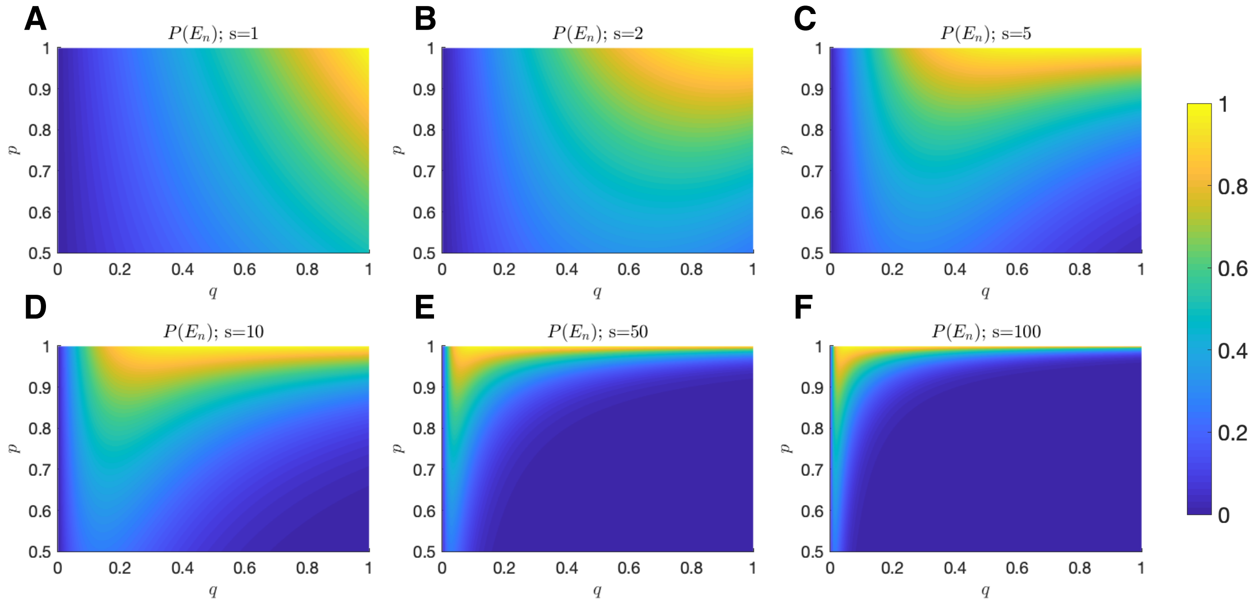


Figure S4: Tie Probability. The probability of a tie is plotted as a function of $q$ (x-axis) and $p$ (y-axis) for (A) $s = 1$; (B) $s = 2$; (C) $s = 5$; (D) $s = 10$; (E) $s = 50$; (F) $s = 100$.

#### S2.3.1 Exact dynamics

Let $I_F$ denote the usual indicator random variable on event $F$:

$$I_{F(\omega)} = \begin{cases} 1, & \omega \in F; \\ 0, & \omega \notin F. \end{cases} \tag{S14}$$

If $r_n$ is unknown, then the distribution of $\ell_n$ follows that of $r_n$ on a strictly positive outcome normalized to the probability of surviving:

$$\begin{aligned}\mathbb{P}(\ell_n = \ell \mid E_n) &= \mathbb{P}([\ell_n = \ell] \cap [\ell_n = r_n > 0]) \,/\, \mathbb{P}(E_n) \\ &= \mathbb{P}(r_n = \ell_n = \ell > 0) \,/\, (\eta^{s_n} - \gamma^{s_n}) \\ &= \begin{cases} m(\ell,\ell)/\,\mathbb{P}(E_n), & 0 < \ell \le s_n; \\ 0, & \ell = 0. \end{cases} \\ &= I_{[\ell>0]} \binom{s_n}{\ell} (pq)^{\ell} (1-q)^{s_n-\ell} / (\eta^{s_n} - \gamma^{s_n}) . \end{aligned} \tag{S15}$$

In this case, the mean number of lost antigens conditioned on a tie becomes

$$\begin{aligned}\mathbb{E}[\ell_n \mid E_n] &= \sum_{\ell=0}^{s_n} \ell\, \mathbb{P}(\ell_n = \ell \mid E_n) \\ &= (\eta^{s_n} - \gamma^{s_n})^{-1} \sum_{\ell=1}^{s_n} \ell \binom{s_n}{\ell} (pq)^{\ell} (1-q)^{s-\ell} \\ &= \frac{pq\eta^{s_n-1}}{\eta^{s_n} - \gamma^{s_n}} s_n. \end{aligned} \tag{S16}$$

Of course, for any realized number of recognized antigens $r_n$ at period $n$ (event $F_{n,r} = [r_n = r]$), the number of lost antigens conditional on a tie $\ell_n$ is completely determined, since

$$\mathbb{P}(\ell_n = \ell \mid E_n \cap F_{n,r}) = \mathbb{P}(\ell_n = \ell \mid \ell_n = r_n = r > 0) = I_{[\ell=r]}, \tag{S17}$$

so that the conditional mean number of lost antigens must match exactly those recognized:

$$\mathbb{E}[\ell_n \mid E_n \cap F_{n,r}] = \sum_{\ell=0}^{s_n} \ell \cdot \mathbb{P}(\ell_n = \ell \mid E_n \cap F_{n,r}) = \sum_{\ell=0}^{s_n} \ell I_{[\ell=r]} = r, \tag{S18}$$

### S2.4 Mean transition behavior

The state transition equation for this process is given by

$$s_{n+1} = s_n - \ell_n + f_n, \tag{S19}$$

where the $\{f_n\}_n$ represent the arrival of new antigens. In our model we will assume that they are IID random penalties with mean $\mathbb{E}[f_n] = f$ and finite variance (e.g. Poisson-distributed, for example). Given this, we will now characterize the mean transition behavior conditioned on a tie and the information available at the present moment. We write $\mathbb{E}_n[\cdot]$ to denote the conditional expectation with respect to date-$n$ information.

#### S2.4.1 Exact dynamics

The mean number of detectable antigens evolves according to the difference equation

$$\begin{aligned}\mathbb{E}_n[s_{n+1} \mid E_n] &= \mathbb{E}_n[s_n - \ell_n + f_n \mid E_n] \\ &= \mathbb{E}_n[s_n] - \mathbb{E}_n[\ell_n \mid E_n] + \mathbb{E}[f_n] \\ &= s_n - \frac{qp\eta^{s_n-1}}{\eta^{s_n} - \gamma^{s_n}} s_n + f, \end{aligned} \tag{S20}$$

which follows since $s_n$ is known at date-$n$ and independent from $E_n$, while $f_n$ is independent from date-$n$ and $E_n$. This process is mean stationary at $s_n = \mu$ whenever

$$\Delta s_n \equiv \mathbb{E}_n[s_{n+1} \mid E_n] - s_n = 0 \;\Rightarrow\; f = \frac{qp\eta^{\mu-1}}{\eta^{\mu} - \gamma^{\mu}} s_n; \tag{S21}$$

or, alternatively,

$$\mu = (f/q) \left( \frac{\eta^{\mu} - \gamma^{\mu}}{p\eta^{\mu-1}} \right). \tag{S22}$$

Plots of fixed points of Eq. S20 are illustrated in Fig. S5 for $p > 1/2$ and $q$ away from zero for small mean penalties $f$. As expected, increases in mean penalty result in higher equilibria. In the large $\mathbb{P}(E_n)$ region of interest, increased $q$ results in a lower number of detectable antigens at equilibrium since more are recognized during each period.

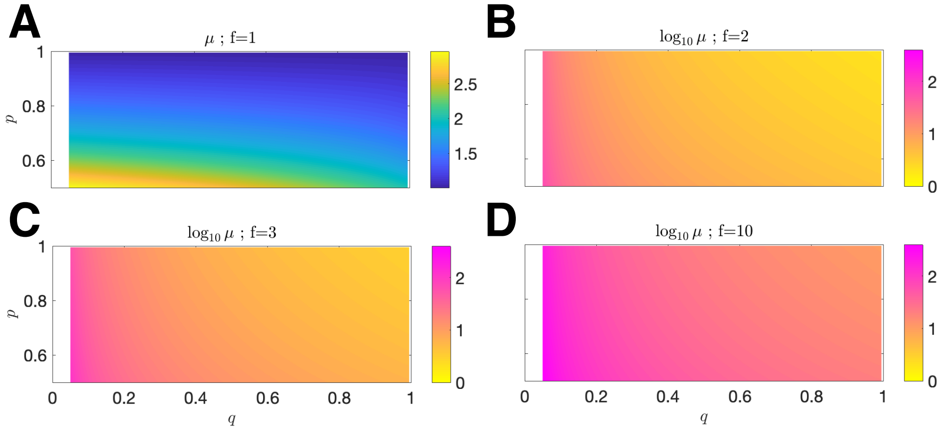


Figure S5: Mean equilibrum. The mean equilibrium point $\mu$ is plotted as a function of $q$ (x-axis) and $p$ (y-axis) for mean penalty (A) $f = 1$; The normalized $\log_{10} \mu$ vs $q$ and $p$ is plotted for mean penalty (B) $f = 2$; (C) $f = 3$; (D) $f = 10$ (In all cases, the relevant domain is restricted to $(p, q) \in (0.5, 1) \times (0.05, 1)$).

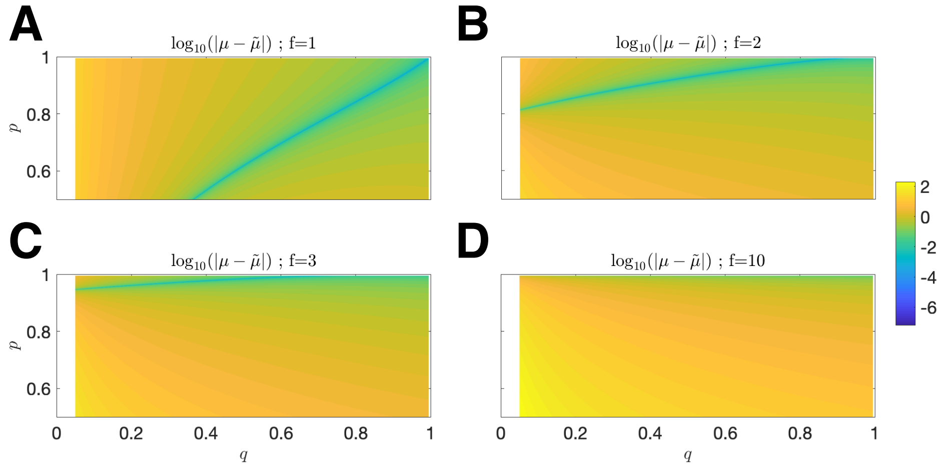


Figure S6: Error of upper estimate $\tilde{\mu}$. The log-absolute error $\log_{10}(|\mu - \tilde{\mu}|)$ vs. $q$ and $p$ is plotted for mean penalty (A) $f = 1$; (B) $f = 2$; (C) $f = 3$; (D) $f = 10$ (In all cases, the relevant domain is restricted to $(p, q) \in (0.5, 1) \times (0.05, 1)$).

#### S2.4.2 Approximate dynamics

If $r_n$ is explicitly given then the mean transition equation simplifies to

$$\begin{aligned}\mathbb{E}_n\left[s_{n+1} \mid E_n \cap F_{n,r}\right] &= \mathbb{E}_n\left[s_n - \ell_n + f_n \mid E_n \cap F_{n,r}\right] \\ &= s_n - \mathbb{E}_n\left[\ell_n \mid E_n \cap F_{n,r}\right] + \mathbb{E}\left[f_n\right] \\ &= s_n - \mathbb{E}_n\left[r_n\right] + f \\ &= s_n - r_n + f, \end{aligned} \tag{S23}$$

since $s_n$ is known at date-$n$, while $f_n$ is independent from date-$n$ and $E_n \cap F_{n,r}$. We can use this to approximate the exact recognition dynamics described above by assuming $r_n = \mathbb{E}_n\left[r_n\right] = qs_n$. In this case,

$$\begin{aligned}\mathbb{E}_n\left[s_{n+1} \mid E_n \cap F_{n,r}\right] &= s_n - qs_n + f \\ &= (1-q)s_n + f. \end{aligned} \tag{S24}$$

The equilibrium may be given explicitly as

$$\tilde{\mu} = f/(1-\gamma) = f/q. \tag{S25}$$

We distinguish the approximate equilibrium $\tilde{\mu}$ from that of exact case $\mu$, the latter incorporating a correction term arising from the fact that knowledge of a tie requires a larger average value of $r_n$ above $qs_n$, since ties occur only when $r_n > 0$. We remark that the equilibria given by Eqs. S22 and S25 are close to one another for small penalty (Fig. S6) and parameter regions that overlap with those having large tie probabilities ($p \sim 1$, $q > 0.5$; Fig. S4), which intuitively suggests that a process driven by its mean overlaps well with one conditional on a tie provided the escape and elimination probabilities are small. We obtain good agreement between averages of large-scale simulations of the process, together with the predicted exact and approximate equilibria for $p, q > 0.5$ and small penalty (Fig. S7). Of course, the the mean dynamics are also approximate since $qs_n$ is in general non-integer-valued. With this in mind, we focus on the dynamics given by Eq. S23.

Here, $r_n$ is Binomially-distributed conditional on the number of current antigens, so that

$$\mathbb{E}_n\left[r_n\right] = qs_n; \qquad \mathbb{V}\mathrm{ar}_n\left[r_n\right] = q(1-q)s_n. \tag{S26}$$

We define the following zero-mean noise variable

$$\varepsilon_n \equiv (f_n - f) - (r_n - qs_n), \tag{S27}$$

and re-write Eq. S19 as

$$s_{n+1} = \gamma s_n + f + \varepsilon_n. \tag{S28}$$

This is none other than a first-order autoregressive, or AR(1), process with innovation terms $\varepsilon_n$ comprised of endogenous noise due to the variance in the number of recognized antigens and exogenous noise generated by fluctuations in the random penalty term.

The process is stable for all but trivial choices of probability $\gamma$. The mean behavior evolves according to:

$$\mathbb{E}_n\left[s_{n+1}\right] = \mathbb{E}_n\left[\gamma s_n + f - \varepsilon_n\right] = \gamma s_n + f, \tag{S29}$$

which implies that

$$\begin{aligned}\mathbb{E}\left[s_n\right] &= \gamma^n s_0 + f\sum_{j=0}^{n-1}\gamma^j \\ &= \gamma^n s_0 + \left(\frac{1-\gamma^n}{1-\gamma}\right)f \\ &\to f/q \ \text{ as } \ n \to \infty, \end{aligned} \tag{S30}$$

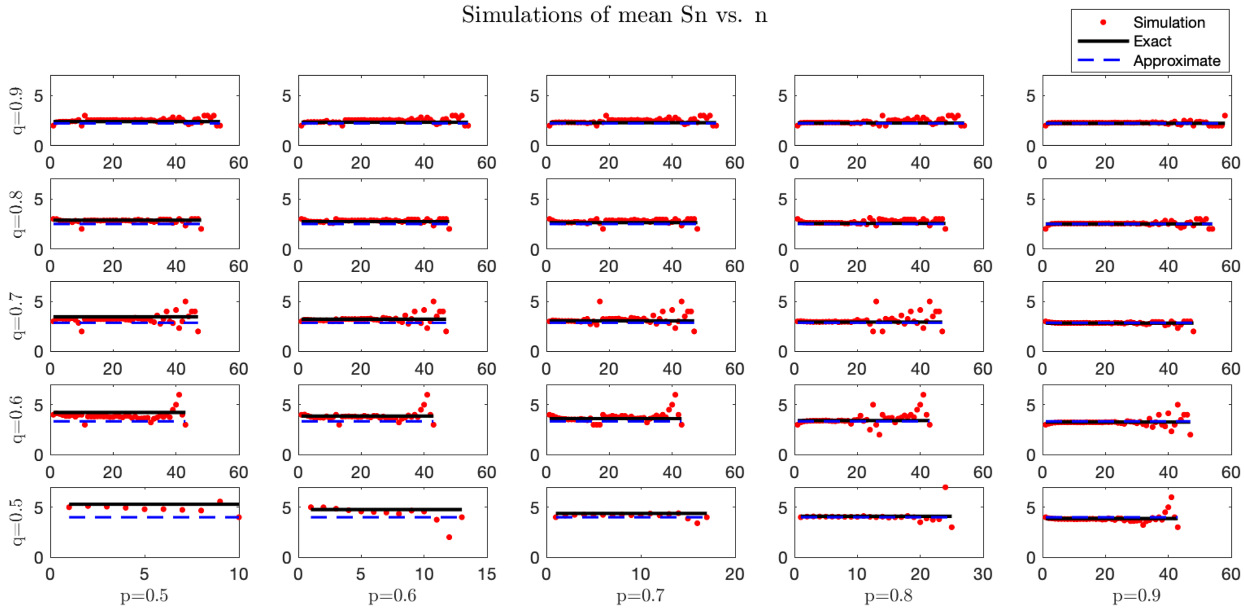


Figure S7: Simulated and analytical mean equilibrium. Stochastic simulations were performed for a process with $s_0$ set equal to the rounded predicted $\mu$. For each trajectory, the pre-escape/elimination time and number of antigens are recorded, and an average is taken for each time. Simulation averages (red) are compared with predicted exact (black solid line) and approximate (blue dashed line) mean dyanmics (For each case, $n = 10^6$ simulations were performed under small mean penalty $f = 2$. Columns from left-to-right correspond with $p \in \{0.5, 0.6, 0.7, 0.8, 0.9\}$. Rows from bottom-to-top correspond with $q \in \{0.5, 0.6, 0.7, 0.8, 0.9\}$.

thus showing agreement in mean with the fixed point given by Eq. S25. Of course, $s_n = \tilde{\mu} = f/q$ satisfies the martingale property ($\mathbb{E}[s_{n+1}] = \gamma f/q + f = f/q = s_n$), and the process tends toward equilibrium with expected inter-temporal difference

$$\Big|\mathbb{E}[s_{n+1}] - \mathbb{E}[s_n]\Big| = \left|(\gamma^{n+1} - \gamma^n)s_0 + f\gamma^n\right| = \gamma^n\left|f - qs_0\right|. \tag{S31}$$

The variance at stationarity, $\mathbb{V}\mathrm{ar}(s_n)$, can be calculated by solving for the fixed point of

$$\mathbb{V}\mathrm{ar}(s) = \gamma^2\mathbb{V}\mathrm{ar}(s) + \sigma_f^2, \tag{S32}$$

giving,

$$\mathbb{V}\mathrm{ar}(s_n) = \sigma_f^2/(1-\gamma^2). \tag{S33}$$

### S2.5 Recognizer Success Probability

For the event $W_n$ (resp. $L_n$) that the Recognizer wins (resp. loses) at period $n$, and for the event $E_n$ of a tie at period $n$, we have,

$$\mathbb{P}(W_n) = \mathbb{P}(E_{n-1})(1-\eta^{s_n}), \tag{S34}$$
$$\mathbb{P}(L_n) = \mathbb{P}(E_{n-1})\gamma^{s_n}, \tag{S35}$$
$$\mathbb{P}(E_n) = \mathbb{P}(E_{n-1})(\eta^{s_n} - \gamma^{s_n}). \tag{S36}$$

These relationships, along with the implicit evolution given by Eq. S25 are used to approximate ultimate Recognizer success probabilities for all possible $p$ and $q$ against several choices of initial antigen number

$s_0$ and mean penalty $f$, and are compared with simulations of using actual transitions via Eq. S21 (Fig. S8). We find good agreement between these methods in characterizing the final outcome over a variety of parameter choices, where accuracy is highest in the relevant parameter region of interest. In particular, the left-column of Fig. S8 details the likelihood that a (static) threat is controlled in the special case where no penalty is assumed.

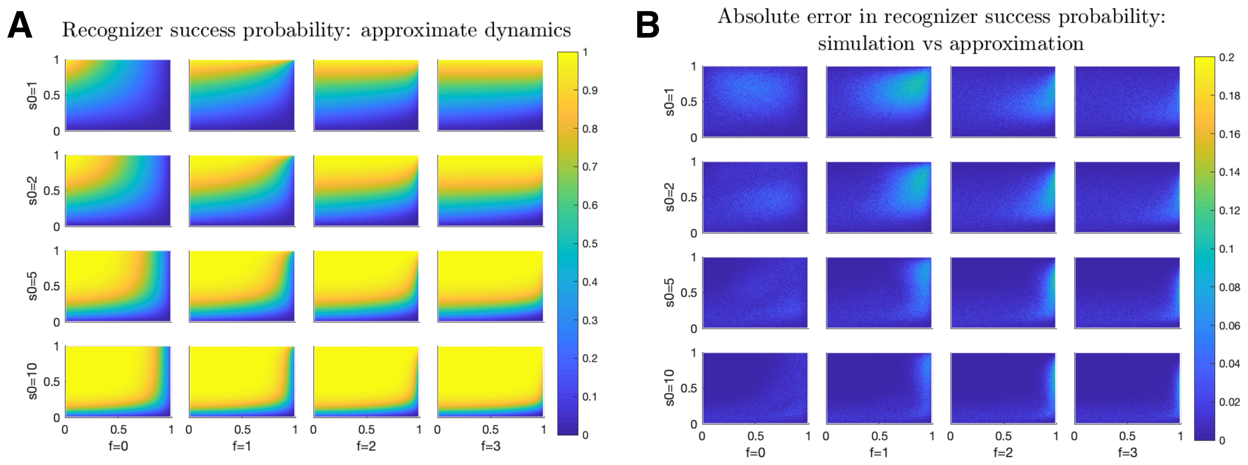


Figure S8: Probability of Recognizer Success. Recognizer success probabilities are plotted as a function of $p$ ($x$-axis) and $q$ ($y$-axis) for (A) approximate dynamics (Eq. S23; (B) The absolute difference between the above approximations and simulated exact dynamics given by Eq. S23. (Results averaged over 1000 iterations for each $(q, p, s_0, f)$ quintuplet. Columns from left-to-right correspond with $f \in \{0, 1, 2, 3\}$. Rows from top-to-bottom correspond with $s_0 \in \{1, 2, 5, 10\}$).

### S2.6 Mutation accumulation rate and tumor antigen availability

The above analysis was motivated by a desire to explain both genetic and non-genetic possibilities leading to recognition evasion. We can consider applying this model to strictly describe genetic evasion in the form of somatic mutations leading either to the generation of (recognizable) tumor-associated antigens or to escape via the removal of these antigens. Using the above framework, mutations, denoted by $\lambda$, accumulate across each period in proportion to the sum of antigens down-regulated to enhance escape and antigens gained via penalty. Thus their rate of accumulation may be expressed by

$$\nu(n) \equiv \frac{\Delta\lambda(n)}{\Delta n} \propto \ell_n + f_n. \tag{S37}$$

Together with the fact that $\ell_n = r_n$ during progression, we have for the mean rate of mutant accumulation

$$\begin{aligned} \mathbb{E}\left[\nu(n)\right] &\propto \mathbb{E}\left[\mathbb{E}\left[r_n \mid s_n\right] + f\right] \\ &= q\mathbb{E}\left[s_n\right] + f \\ &\to 2f \quad \text{as} \quad n \to \infty, \end{aligned} \tag{S38}$$

ultimately giving

$$\lambda(n) \propto 2fn. \tag{S39}$$

which predicts that the rate of mutational acquisition is linear in time, consistent with empirical observation [1, 2]. Heuristically, tumors that survive while accumulating an average of $f$ targetable alterations must balance those gains by $f$ additional evasion events. This theory predicts, perhaps surprisingly, that the mutation rate is a direct reflection of the penalty paid for cancer progression as a function of its local environment. Tumors having a more difficult time surviving in a hostile or restrictive environment would be predicted to have higher rates of mutation. In this context, high mutational signatures are predicted to be correlated with tumors that are more susceptible to recognition. For a passive Evader, our theory predicts that the observed mutation rate depends only on the mean penalty term for cancer progression, unaffected by recognition rate. On the other hand, the stationary number of available antigens, approximated by $\tilde{\mu} = f/q$, varies directly with evasion penalty and inversely with antigen recognition rate. Moreover, mutation or adaptation accumulation is expected to converge to a stable equilibrium for any recognition, evasion, and penalty rates.

### S3 Active Evader in an Adaptive Environment

In the previous section we considered the predicted dynamical behavior when the Evader is assumed to adopt a fixed strategy. In that case, if number of detectable antigens is moderately large ($s_0 \sim 10$), then the game is biased against the Evader for most combinations of evasion and recognition success probabilities (Sec. S2.2). Additionally, mean transitions in the number of recognizable antigens obey an AR(1) process tending toward the quotient of the mean penalty and recognition rate (Sec. S2.4). Moreover, this behavior predicts that the observed mutation accumulation rate is linear in time and proportional to the mean penalty term (Sec. S2.6). Here, we allow for the Evader to optimally select his evasion rate at each period. Larger success rates come at the cost of adding back more recognition opportunities in the subsequent time step, so that the Evader employs a strategy to maximize his survival or likelihood of escape. This framework is motivated by the observation that cancer threats are known to accumulate perhaps mildly deleterious mutations that occur passively during evolution to obtain rare 'driver' mutations [3]. The novelty here is that we propose a unifying theoretical framework to investigate the resulting strategy employed by a cancer population if the choice of evasion is planned based on knowledge of the current antigen landscape and hostility, or number of recognized targets.

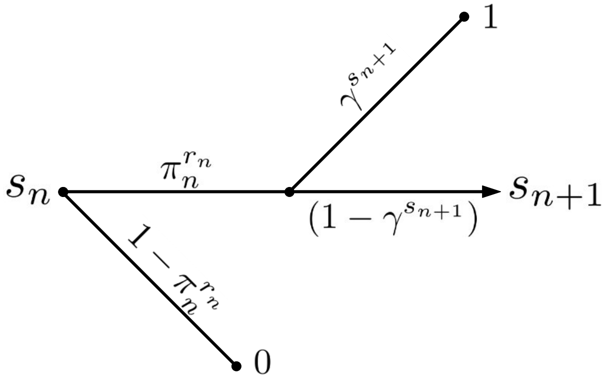


Figure S9: Active Evader Decision Tree. This process is identical to that of Fig. S2, except that in this case the evasion rate $\pi_n$ may be chosen at each period by the evader.

In contrast with the prior section, which considered temporal evolution as a function of fixed evasion rate $p$ and random penalty $f_n$, here, the evasion rate $\pi_n$ may depend on time, and for simplicity we consider deterministic penalties. In order to properly frame this problem in a manner suitable to handle via dynamic programming, we define the necessary parameters, expectation, and value functions below. We assume that the process evolves according to state transition equation,

$$s_{n+1} = s_n - r_n + f_n, \tag{S40}$$

and that conditional expectations are taken with respect to $\mathcal{F}_n$ (the natural filtration [4] with respect to the underlying process). If at time $n$, knowledge of total $s_n$ and recognized $r_n$ targets is known, than the Evader's objective is to select a policy $\pi \equiv \{\pi_n, \pi_{n+1}, \dots\}$ that maximizes the sum of present and future rewards, $R(s_n, r_n, \pi_n)$, which in general depend on the current state, $s_n$, as well as the Recognizer, $r_n$, and Evader, $\pi_n$, actions. The value function is defined to be the maximal attainable sum of expected future rewards, given by

$$J_n(s_n) = \sup_{\pi} \mathbb{E}_n \left[ \sum_{m=n}^{\infty} R\left(s_m, r_m, \pi_m\right) \right]. \tag{S41}$$

Problems which may be framed in this context have been well-studied and utilize a rich theory of stochastic dynamic programming, originally proposed by Richard Bellman [5,6]. Bellman's Principal of Optimality and

Bellman equation for a stationary solution (independent of starting time) is given via backward induction by

$$J(s_n) = R(s_n, r_n, \pi_n) + J(s_{n+1}). \tag{S42}$$

Eq. S42 states that the maximal attainable value at period $n$ is given by the sum of the maximal attainable value at the next time step, $J(s_{n+1})$, and the $n$-period reward of strategy $\pi_n$ obeying Eq. S41. For the

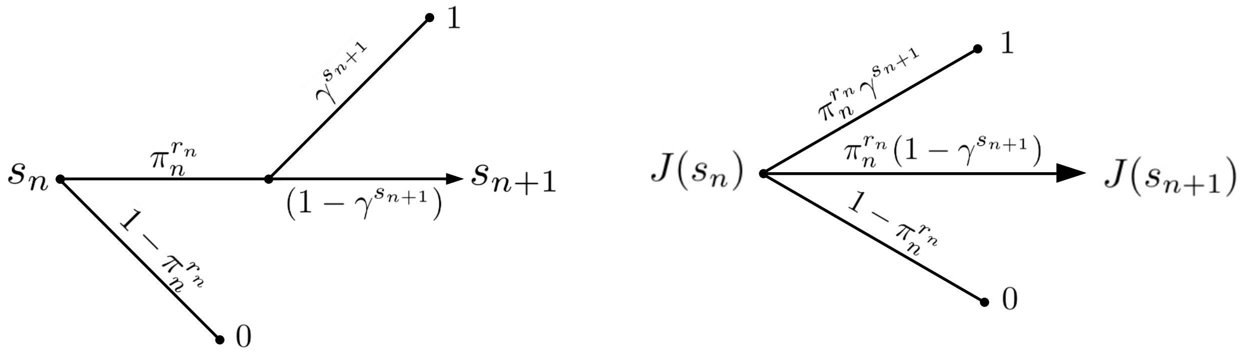


Figure S10: Active Evader Decision Tree. The value function for the Evader $J$ is represented under stationary dynamics. The reward of Evader elimination is 0, and for successful escape is assumed to be 1 with no time-additive discounting.

problem at hand, we assume that the Evader receives a normalized reward of either $R_n = 1$ if it escapes at any time period (there is no temporal discount for escape at later periods), or $R_n = 0$ if it is eliminated. In this case, we may draw a decision tree for the $n$-period problem in terms of the value function $J$, current antigen number $s_n$, Recognizer antigen recognition miss probability $\gamma = 1-q$, number of recognized antigens $r_n$, and Evader strategy, $\pi_n$ (Fig. S10). Here, $\pi_n$ represents the $n$-period probability of antigen loss by the Evader.

Using the dynamic programming principle, the Bellman equation under uncertainty takes the form

$$J\big(s_n\big) = \max_{\pi_n} \Big\{ \mathbb{E}_n \Big[ \pi_n^{r_n} \big[ \gamma^{s_{n+1}} + \big(1 - \gamma^{s_{n+1}}\big) J\big(s_{n+1}\big) \big] \Big] \Big\}. \tag{S43}$$

Under a particular choice of assumed penalty and transition equation, we can calculate an exact, closed-form solution to the dynamic program in Eq. S43. This solution generates an optimal policy, given by $\pi^* = \{\pi_1^*, \pi_2^*, \ldots, \pi_n^*, \ldots\}$, a sequence of optimal decisions, in addition to the maximal value at each time assuming the optimal policy, given by $J(s_n)|_{\pi_n^*}$.

### S3.1 Constitutive relations for inter-temporal penalty

We make the following assumptions in our setting to make this problem more tractable. The first assumption is that the penalty function is time-homogeneous and deterministic:

$$f(s_n, r_n, \pi_n), \quad \pi_n \in [0,1], \quad s_n, r_n \in \mathbb{Z}^+. \tag{S44}$$

Conditional on progressing to the next period, the transition equation takes the following form:

$$s_{n+1} = s_n - r_n + f(s_n, r_n, \pi_n). \tag{S45}$$

In cases where we wish to emphasize the dependence of the transition equation on $\pi_n$, we will denote $s_{n+1}$ by $g(\pi_n)$ so that

$$g(s_n, \pi_n) = s_{n+1}. \tag{S46}$$

The second assumption is that this penalty is $\pi_n$-affine. That is,

$$f(s_n, r_n, \pi_n) = h_m(s_n, r_n)\pi_n + h_b(s_n, r_n). \tag{S47}$$

for positive $h_m$ and $h_b$. We also assume that the baseline penalty is independent of $s_n$ so that $h_b = h_b(r_n)$.

In order to analytically characterize the solution we assume that $r_n$ is known prior to choosing $\pi_n$ ($r_n \in \mathcal{F}_n$). In the analogous coin game, the Evader is allowed to see the success of his opponent, the Recognizer, prior to choosing a strategy. In this case, the dynamic program has a solution if we also assume that the linear penalty term can be represented by

$$h_m(s_n, r_n) = \frac{r_n}{c}\left(\frac{1}{\delta_n} \cdot \frac{1-\gamma^{s_n}}{1-\gamma}\right)^{1/r_n} \tag{S48}$$

with $c \equiv -\ln\gamma > 0$ and $0 < \delta_n \leq 1$. This assumption implies that the marginal penalty of increasing $\pi_n$ is asymptotically proportional to the number of recognized antigens. This is reasonable to assume, for example, in cases where significant immune system recognition and tumor killing create an environment that makes subsequent adaptation more costly, resulting possibly from increased inflammation. The constant $\delta_n$, a free variable, is inversely related to aversion of the Evader strategy so that larger values imply a bolder evasion strategy for all else held constant. This parameter may in general vary temporally and as a function of disease sub-type.

### S3.2 Dynamic programming solution

In the above case, we may find an exact solution to the optimal programming problem. Since $r_n \in \mathcal{F}_n$ (the filtration generated by the evolution of $s_n$ and the Recognizer action at time $n$), the stationary Bellman Equation takes the form

$$J\big(s_n\big) = \max_{0\leq\pi_n\leq 1}\Big\{\pi_n^{r_n}\big[\gamma^{s_{n+1}} + \big(1-\gamma^{s_{n+1}}\big)J\big(s_{n+1}\big)\big]\Big\}. \tag{S49}$$

For simplicity in the subsequent definition we drop the period index, re-writing Eq. S49 as

$$J\big(s\big) = \max_{0\leq\pi\leq 1}\Big\{\pi^{r}\big[\gamma^{g(s,\pi)} + \big(1-\gamma^{g(s,\pi)}\big)J\big(g(s,\pi)\big)\big]\Big\} \tag{S50}$$

Using $c \equiv -\ln\gamma$, the first-order condition (FOC) is

$$\frac{\partial}{\partial\pi}\Big\{\pi^{r}\big[e^{-cg(s,\pi)} + \big(1-e^{-cg(s,\pi)}\big)J\big(g(s,\pi)\big)\big]\Big\} = 0. \tag{S51}$$

In expanded form, the FOC becomes

$$\pi^{r-1}\Bigg\{r\Big[e^{-cg} + \big(1-e^{-cg}\big)J(g)\Big] + \pi\Big[-c\frac{\partial g}{\partial\pi}e^{-cg} + c\frac{\partial g}{\partial\pi}e^{-cg}J(g) + (1-e^{-cg})\frac{\partial J}{\partial g}\frac{\partial g}{\partial\pi}\Big]\Bigg\} = 0. \tag{S52}$$

From Eq. S47, we have that

$$\frac{\partial f}{\partial\pi} = \frac{\partial g}{\partial\pi} = h_m. \tag{S53}$$

We postulate that the solution takes the following form:

$$J(s) = \frac{A\gamma^s}{1-\gamma^s}. \tag{S54}$$

so that

$$\frac{\partial J}{\partial s} = -\frac{cJ(s)}{(1-e^{-cs})}. \tag{S55}$$

This, together with Eq. S53 reduces Eq. S52 to

$$\pi^{r-1}\Big[e^{-cg} + \big(1-e^{-cg}\big)J(g)\Big]\big(r - ch_m\pi\big) = 0. \tag{S56}$$

Thus, the optimal Evader success probability, $\pi^*$, is given by

$$\pi^* = r/ch_m. \tag{S57}$$

Under Evader optimal strategy, the transition equation in Eq. S45 becomes

$$g^* \equiv g(s,\pi^*) = s - r + f(s,r,\pi_*) = s - r + \big(r/c + h_b\big). \tag{S58}$$

We next confirm that this satisfies the Bellman Equation (Eq. S50). The above solution implies

$$J\big(s\big) = \pi^{*r}\big[\gamma^{g^*} + \big(1-\gamma^{g^*}\big)J\big(g^*\big)\big], \tag{S59}$$

which ultimately yields

$$A\gamma^s = \delta(1-\gamma)(1+A)\gamma^{h_b+r/c-r}\gamma^s. \tag{S60}$$

Equating coefficients and applying this logic to each policy gives

$$A_n = \frac{\delta_n(1-\gamma)\gamma^{h_b+r/c-r}}{1-\delta_n(1-\gamma)\gamma^{h_b+r/c-r}}. \tag{S61}$$

The optimal policy is given by the sequence

$$\pi_n^* = \left(\frac{\delta_n(1-\gamma)}{1-\gamma^{s_n}}\right)^{1/r_n}. \tag{S62}$$

We henceforth refer to $\delta_n$ as the aversion parameter. Large values of $\delta_n$ imply low aversion. It can be interpreted as the selected strategy in the simplest case where $\delta_n = \delta > 0$ and $s_n = r_n = 1$, since

$$\pi_n^* = \delta; \tag{S63}$$

Rearranging Eq. S62 gives

$$\frac{1-\gamma_n^s}{1-\gamma} = \frac{\delta}{\pi_n^{*\,r_n}}. \tag{S64}$$

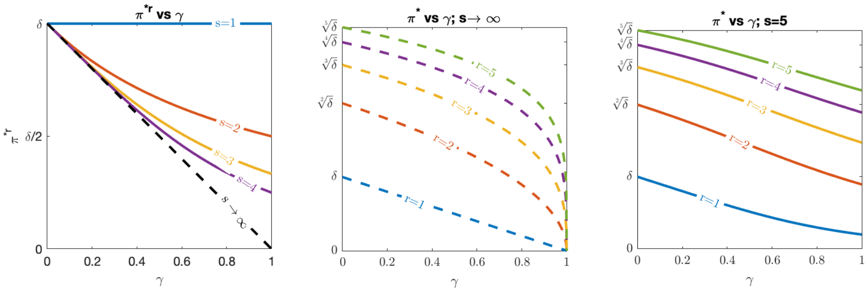


Figure S11: Optimal evasion. (Left) $\pi^* r$ vs $\gamma$, (Middle) $\pi^*$ vs. $\gamma$ for $s \to \infty$, and (Right) $\pi^*$ vs. $\gamma$ for $s = 5$ under various values of recognized antigens, assuming $\delta < 1$.

### S3.3 Solution uniqueness

**Proposition.** *The above value function is unique.*

*Proof.* We consider value functions $V(s)$ in the space of functions that are continuous in $\pi$ and bounded in $s$. We take $||V||_\infty \equiv \sup_s |V(s)|$. From the previous section, we have identified such a function $J$ so that

$$J(s_n) = \max_{0\leq\pi\leq 1} \pi^r \left[\gamma^{s_{n+1}} + (1-\gamma^{s_{n+1}})J(s_{n+1})\right]. \tag{S65}$$

Assume that $V(s)$ is another solution. For fixed $s_n$, let $\pi^*$ be such that

$$V(s_n) = \pi^{*r} \left[\gamma^{s_{n+1}} + (1-\gamma^{s_{n+1}})V(s_{n+1})\right]. \tag{S66}$$

We can re-write the following term:

$$\gamma^{s_{n+1}} = \gamma^{s_n - r + h_m\pi + h_b} = \gamma^{s_n - r + h_b}(\gamma^{h_m})^\pi \equiv \gamma^{k_s}\tilde{\gamma}^\pi, \tag{S67}$$

where $\tilde{\gamma}, \gamma^{k_s} < 1$. Then

$$\begin{aligned}
V - J &= \pi^{*r}\left[\gamma^{k_s}\tilde{\gamma}^{\pi^*} + (1-\gamma^{k_s}\tilde{\gamma}^{\pi^*})V(s_{n+1})\right] - \max_{0\leq\pi\leq 1}\pi^r\left[\gamma^{k_s}\tilde{\gamma}^\pi + (1-\gamma^{k_s}\tilde{\gamma}^\pi)J(s_{n+1})\right] && \text{(S68)}\\
&\leq \pi^{*r}\left[\gamma^{k_s}\tilde{\gamma}^{\pi^*} + (1-\gamma^{k_s}\tilde{\gamma}^{\pi^*})V(s_{n+1})\right] - \pi^{*r}\left[\gamma^{k_s}\tilde{\gamma}^{\pi^*} + (1-\gamma^{k_s}\tilde{\gamma}^{\pi^*})J(s_{n+1})\right] && \text{(S69)}\\
&= \pi^{*r}(1-\gamma^{k_s}\tilde{\gamma}^{\pi^*})(V(s_{n+1}) - J(s_{n+1}) && \text{(S70)}\\
&\leq \pi^{*r}(1-\gamma^{k_s}\tilde{\gamma}^{\pi^*})\Big|V(s_{n+1}) - J(s_{n+1})\Big| && \text{(S71)}\\
&\leq \pi^{*r}(1-\gamma^{k_s}\tilde{\gamma}^{\pi^*})||V-J||_\infty. && \text{(S72)}
\end{aligned}$$

Note that

$$c(\pi) \equiv \pi^r(1-\gamma^k\tilde{\gamma}^\pi) \leq 1 - \gamma^k\tilde{\gamma}^\pi \tag{S73}$$

is increasing in $\pi$ (since $\tilde{\gamma} < 1$) so that $c(\pi) \leq 1 - \gamma^k\tilde{\gamma} \equiv K < 1$. Thus,

$$V - J \leq K||V-J||_\infty. \tag{S74}$$

By identical argument above, this time reversing the roles of $V$ and $J$ gives

$$J - V \leq K||V-J||_\infty, \tag{S75}$$

and so

$$\Big|V(s_n) - J(s_n)\Big| \leq K||V-J||_\infty < ||V-J||_\infty \text{ for all } s_n. \tag{S76}$$

Therefore,

$$||V-J||_\infty = \sup_{s_n}|V(s_n) - J(s_n)| < ||V-J||_\infty. \tag{S77}$$

Thus,

$$||V-J||_\infty = 0. \tag{S78}$$

□

### S3.4 Mean optimal transitions

From Eq. S58, the mean optimal transitions are:

$$\mathbb{E}_n [s_{n+1} \mid E_n] = s_n + (1/c - 1)r_n + h_b. \tag{S79}$$

The mean increment, $\Delta s_n$, assuming the process is driven by $r_n \sim \text{Binomial}(s_n, q)$, becomes

$$\Delta s_n = (1/c - 1)qs_n + h_b. \tag{S80}$$

We next consider two cases. In the first case, the $\pi$-independent penalty term $h_b$ scales linearly with the number of currently recognized antigens, and in the second case this independent penalty term is assumed fixed.

#### S3.4.1 $\mathbf{r_n}$-linear $\pi_\mathbf{n}$-affine penalty

This case considers $h_b = \alpha r_n$. Here, larger recognition in the current period results in larger exogenous penalty, and hence easier targeting, in the next period. Consequently, the number of detectable antigens in the future is directly influenced by both the tumor evasion strategy $\pi^*$ and the extent of that recognition resulting from immune targeting $r_n$. In this case, we have that

$$\mathbb{E}[(1/c - 1 + \alpha)r_n \mid s_n] = (1/c - 1 + \alpha)qs_n, \tag{S81}$$

so that the process satisfies the Martingale condition

$$\mathbb{E}[s_{n+1} \mid s_n] = s_n \tag{S82}$$

for critical alpha

$$\alpha_c = \frac{\log \gamma^{-1} - 1}{\log \gamma^{-1}}. \tag{S83}$$

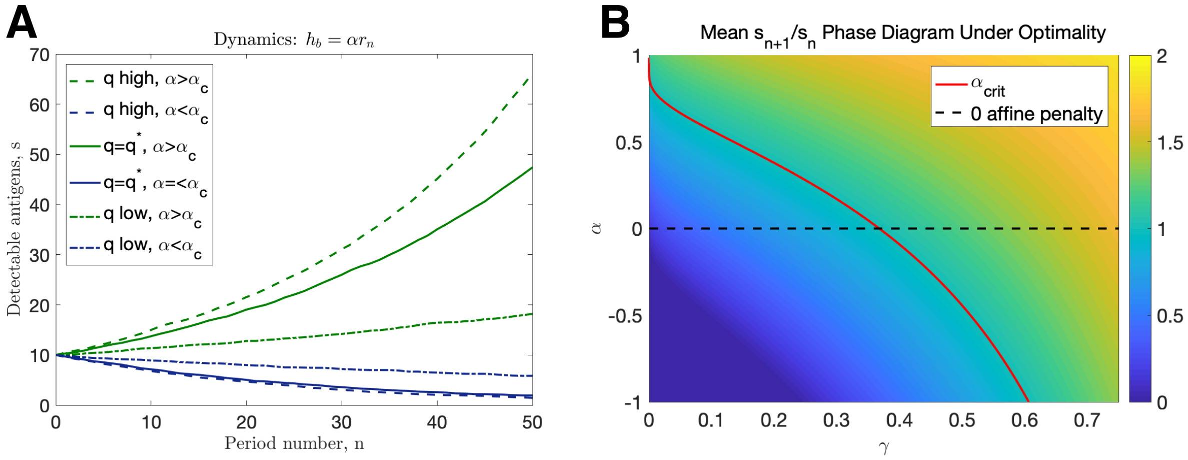


Figure S12: $r_n$-linear dynamics. (A) Simulated trajectories assuming $h_b = \alpha r_n$ ($q$ high: $q = 0.75$, $q^* = 1-1/e$, $q$ low: $q = 0.25$. $\alpha > \alpha_c$: $\alpha = \alpha_c+0.05$, $\alpha_c$ given by Eq. S83, $\alpha < \alpha_c$: $\alpha = \alpha_c-0.05$); (B) Mean intertemporal gain as a function of $\gamma = 1 - q$ and $\alpha$.

**Mutation accumulation rate** In the trivial case where, $\alpha = \alpha_c$, $s$ is constant and so mutation accumulation is predicted to be linear. Contributions by optimal evasion to the mutation rate are expected to exponentially decrease (resp. increase) over time if $\alpha < \alpha_c$ (resp. $\alpha > \alpha_c$).

In this case, dynamics and resultant mutation accumulation is determined by $\alpha$ relative to $\alpha_c$, and only those $\alpha$ close to the threshold generate behavior resembling linear mutation accumulation. Given this, the added penalty $h_b(r_n) = \alpha r_n$ due to the number of recognized antigens appears to be a less reasonable assumption based on empirical mutation rates [1, 2]. We next consider the $r$-independent $\pi_n$-affine penalty case $h_b = \beta$.

#### S3.4.2 $\mathbf{r_n}$-independent $\pi_\mathbf{n}$-affine penalty

If recognition incurs no additional penalty on the present evasion strategy, then $h_b = \beta$. In this case, $\Delta s_n$ from Eq. S80 becomes

$$\Delta s_n = (1/c - 1)qs_n + \beta. \tag{S84}$$

The recognition dynamics of this case are more complex and partition into three regimes based on recognition relative to a critical threshold $q^* = 1 - 1/e$ (for which $c = 1$ and Eq. S84 $\Delta s_n = \beta$): Effective immune recognition, critical recognition, and impaired recognition.

**Effective immune recognition:** Here, $q > q^*$, giving $c > 1$. In this case, the Recognizer exerts a large recognition rate on the evading tumor. If $h_b = \beta \leq 0$, then the equilibrium, $s^*$ for which $\Delta s_n = 0$ is negative, and the $s_n$ is driven to 0. If $h_b$ is a positive penalty against the Recognizer, then there exists a stable, positive antigen state:

$$s^* = \frac{\beta}{q(1 - 1/c)} \tag{S85}$$

Trajectories assuming a variety of initial conditions are given with $s^* = 10$ in Fig. S13A.

**Impaired immune recognition:** In contrast with effective recognition $q < q^*$, $c < 1$, and in this case, the equilibrium points are unstable. Moreover, If $h_b = \beta \geq 0$, then by a similar reasoning as above, $s^* \leq 0$ so that $s_n$ is driven to become very large. Alternatively, if $h_b < 0$ then the equilibrium state is:

$$s^* = \frac{\beta}{q(1/c - 1)} \tag{S86}$$

**Critical immune recognition:** At criticality $q = q^*$, $c = 1$, and Eq. S80 simplifies to

$$\Delta s_n = \beta. \tag{S87}$$

In this special case, all randomness imparted to the process by $r_n$ is eliminated by a critical offset in the number of recognized antigens and the penalty so that the long-term behavior of the process is completely determined by $h_b = \beta$. Predictably, $\beta > 0$ (resp. $\beta < 0$) results in net expansion (resp. depletion) of antigens over time, and $\beta = 0$ is stationary. The sign of $\beta$ may change as a function of the tumor IME. For example, immune exclusion and the resulting attenuated inflammation may both decrease $q$ and $\beta$ as well as genetic aberrations involving mismatch repair (MMR) deficiency and microsatellite instability. Other alterations, such as modulated MHC expression, or MHC loss of heterozygosity (LOH), may affect $q$ in isolation [7].

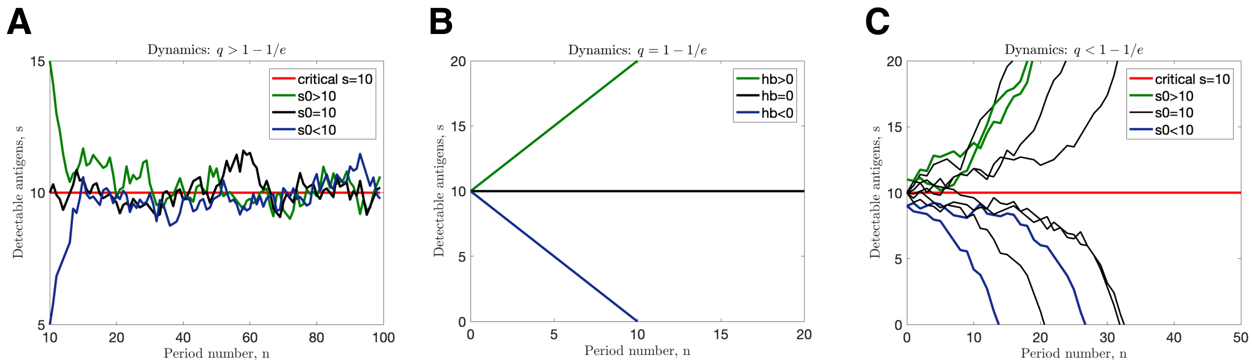


Figure S13: Transition dynamics. (A) $q > q^*$ and $\beta = 1.76$; (B) $q = q^*$; (C) $q < q^*$ and $\beta = -1.68$ (in all cases illustrated, $\beta$ chosen so that $s^* = 10$).

**Mutation accumulation rate** Critical and impaired immune recognition dynamics follow a similar behavior to that detailed in Sec. S3.4.1. The effective recognition case bears a resemblance to the approximate dynamics of the informed Evader in Sec. S2.4.2. Here, by a similar argument in Sec. S2.6 once equilibrium is achieved, we have that

$$\nu(n) \equiv \frac{\Delta\lambda(n)}{\Delta n} \propto r_n + f_n. \tag{S88}$$

Studying the process at $s_0 = s^*$ given by Eq. S85, and $f_n^* = r_n/c + \beta$, we have that

$$\begin{aligned}\mathbb{E}_n\left[\nu(n) \mid E_n\right] &\propto \mathbb{E}_n\left[r_n + r_n/c + \beta \mid E_n\right] \\ &= (1 + 1/c)\mathbb{E}_n\left[r_n \mid E_n\right] + \beta \\ &= (1 + 1/c)qs^* + \beta \\ &= \beta\left[\frac{(1 + 1/c)}{(1 - 1/c)} + 1\right] \\ &= \left(\frac{2c}{c - 1}\right)\beta. \end{aligned} \tag{S89}$$

This implies that

$$\lambda(n) \propto 2\beta cn/(c - 1). \tag{S90}$$

Therefore, linear mutation accumulation as a function of time ensues for an effective Recognizer as in the passive Evader case (Eq. S39), this time as a function not only of exogenous affine penalty term $\beta > 0$ but also of $q$ through $c$. We recall that under effective recognition, $q^* < q < 1$ (equivalently $1 < c < \infty$), which ultimately gives via Eq. S90

$$2\beta n < \mu(n). \tag{S91}$$

**Dynamics summary** The assumption of a penalty $f_n = h_m\pi_n + \alpha r_n$ results in exponential growth or decay in the number of recognizable antigens (and therefore mutation rate), and it was only for a very narrow parameter value $\alpha \sim \alpha_c$ for which linear mutation accumulation could occur. It is for this reason that the $r_n$-linear constitutive assumption is less realistic.

For penalties $f_n = h_m\pi_n + \beta$ that are $r_n$-independent, mutations are predicted to accumulate linearly under effective immune recognition, in a similar manner to that observed in the passive Evader case. In contrast with that case however, an active Evader executes an optimal strategy to maximize the overall escape probability. This predicts that one effect of a dynamic evasion that optimally maximizes escape probability is a concomitant increase in the mutation accumulation rate relative to the passive case via a correction term $c/(c-1)$. This enhancement becomes indistinguishable when recognition is very aggressive ($q \to 1$) and becomes large when $q$ approaches the critical detection rate.

Interestingly, the active evasion strategy predicts that mutation accumulation rates vary as a function of recognition pressure, in contrast with the passive evasion model. Additionally, disease progression may affect immune recognition (changes in $q$) and tumor evasion penalty (changes in $\beta$). While the number of recognizable TAAs for the passive case continues evolve according to the mean-reverting process, there is a dramatic discontinuity in active systems whereby recognition rates below a critical threshold may result in unstable behavior prior to escape (Fig. S13).

### S3.5 Optimal evasion strategy

From Eqs. S54, S61, and Eq. S62, we have

$$J(s_n, r_n) = \frac{A_n e^{-cs_n}}{1-e^{-cs_n}}, \tag{S92}$$

$$\tag{S93}$$

$$A_n = \frac{\delta_n q e^{-(1-c)r_n - c\beta}}{1-\delta_n q e^{-(1-c)r_n - c\beta}}, \tag{S94}$$

and

$$\pi_n^* = \left(\frac{\delta_n q}{1-(1-q)^{s_n}}\right)^{1/r_n}. \tag{S95}$$

Thus,

$$J(s_0, r_0) = \frac{\delta_0 q e^{-(1-c)r_0 - c\beta}}{1-\delta_0 q e^{-(1-c)r_0 - c\beta}} \cdot \frac{e^{-cs_0}}{1-e^{-cs_0}}, \tag{S96}$$

We note that for $s_n = s_{n-1} + (1/c-1)r_{n-1} + \beta$, therefore

$$e^{-cs_n} = \gamma^{s_{n-1}+(1-c)r_{n-1}/c+\beta} = \gamma^{s_{n-2}+(1-c)(r_{n-1}+r_{n-2})/c+2\beta} = ... = \gamma^{s_0+n\beta+\mathbb{C}_\gamma R_{n-1}}, \tag{S97}$$

where

$$\mathbb{C}_\gamma \equiv \frac{1-\ln\gamma^{-1}}{\ln\gamma^{-1}} \tag{S98}$$

and

$$R_n \equiv \sum_{j=1}^{n} r_j. \tag{S99}$$

By iteratively applying Eq. S97, we ultimately obtain the value function in terms of the history of the environmental landscape, $\{r_n\}_n$

$$J(s_n, r_n) = \frac{\delta_n q(1-q)^{\mathbb{C}_\gamma r_n - \beta}}{1-\delta_n q(1-q)^{\mathbb{C}_\gamma r_n - \beta}} \cdot \frac{(1-q)^{s_0 - n\beta + \mathbb{C}_\gamma R_{n-1}}}{1-(1-q)^{s_0 - n\beta + \mathbb{C}_\gamma R_{n-1}}}. \tag{S100}$$

We remark that this simplifies for constant $\delta_n = \delta$, which we will typically take as 1.

**Critical recognition:** At the critical value of recognition $q^* = 1 - 1/e$ ($c = 1$), the dynamics become deterministic,. Here, the value of the present state depends only on the initial number of detectable antigens and number of periods that have elapsed and is independent of the history of recognized antigens $\{r_n\}_n$.

$$J(s_n, r_n) = \frac{\delta_n q(1-q)^\beta}{1-\delta_n(1-q)^\beta} \cdot \frac{(1-q)^{s_0 - n\beta}}{1-(1-q)^{s_0-n\beta}}. \tag{S101}$$

At criticality, the value of the present state depends only on the initial number of detectable antigens and number of periods that have elapsed, and not on the number of recognized antigens.

**Non-critical recognition** We recall that the value function carries meaning as the maximal attainable expected future value. Under effective recognition ($c = 1 \Rightarrow \gamma^{\mathbb{C}r}$ is increasing in $r$), so that the value function (Eq. S100) has an exponent that increases

We are motivated to consider either mild or aggressive recognition of Sec. S3.4. We will assume that there is minimal aversion so that $\delta_n = 1$.

#### S3.5.1 Predicted dynamical behavior

From Sec. S3.4.2, the dynamical behavior of the number of recognizable TAAs, or immunogenicity, of an active Evader is determined by $\beta$ and $q$. Disease progression may ultimately affect immune recognition (reducing $q$) and/or tumor evasion penalty (reducing $\beta$). $\beta$ is expected to vary widely across tumor types. Within a given tumor subtype, the extent of environmental hostility is expected to require additional tumor adaptation that may manifest as additional TAA targets. Therefore, larger (resp. smaller) evasion penalties $\beta$ correspond with *anti-tumor* (resp. *pro-tumor*) IME. Similarly, larger (resp. smaller) $q$ corresponds to *infiltrated* (resp. *excluded*) environments, and from this we model four possible states: anti-tumor infiltrated, anti-tumor excluded, pro-tumor infiltrated, and pro-tumor excluded. The model predicts that infiltrated ($q > q^*$) environments lead to an absorbing equilibrium state in the intervening period prior to escape, while exclusion ($q < q^*$) result in unstable equilibria. Interestingly, the sign of the equilibrium, and hence the behavior, depends on $\beta$, and leads to dramatically diverse behavior in the antigenicity of a dominant tumor clone as it progresses via immune recognition. This case is meaningful as long as the inter-temporal penalty assuming the optimal strategy occurs, $f_n^*$, remains non-negative whenever there is at least one recognition event. This is equivalent to the condition that $f_n^* \geq 1/\ln\gamma^{-1} + \beta > 0$, which is assumed in all examples that follow. These results are summarized in Fig. 5 and organized below. The corresponding immunogenicity and cumulative mutations following escape are given by Fig. 4, with the timing of escape and example trajectories given by Fig. S14.

**1. Anti-tumor infiltrated ($q > q^*$, $\beta > 0$):** This stable steady state is positive, so that the process is mean-reverting, and generates immunogenically warm' tumors.

**2. Anti-tumor excluded ($q < q^*$, $\beta > 0$):** Here, recognition is low, while the arrival of new TAAs is large. This unstable steady state is negative, so that all trajectories tend to increase their immunogenicity over time, resulting in 'hot' tumors.

**3. Pro-tumor infiltrated ($q > q^*$, $\beta < 0$):** In this case, recognition is large while the arrival of new TAAs is low. This stable steady state is negative, so that all trajectories tend to reduce their immunogenicity to zero over time, yielding 'cold' tumors.

**4. Pro-tumor excluded ($q < q^*$, $\beta < 0$):** Lastly, if both recognition and new TAA arrival rates are low, then there is a positive unstable state, above which trajectories accumulate additional TAAs over time, becoming 'hot', and below which the populations are predicted to reduce the number of recognizable TAAs over time, becoming 'cold'.

These predicted dynamics parallels the observation that tumors under active immunosurveillance via effective recognition undergo significant immunoediting. Our results predict that the resulting tumor becomes 'warm' or 'cold', depending on the extent of new TAA arrival during active evasion. On the other hand, impaired recognition leads to diverse behavior dependent on the rate at which new TAAs are acquired during active evasion. If this acquisition rate is large, then the tumor accumulates TAAs over time to become 'hot'. On the other hand, tumors subject to reduced selection pressures may evolve as immune-hot or immune-cold tumors, consistent with previous observations [8]. Moreover, the effect of reducing immune recognition leads to an accumulation of TAAs over time, consistent with experimental observations in lung cancer wherein patients with HLA loss of heterozygosity harbored larger mutational burdens, an indirect measure of TAA number of our model [9]. Our predictions suggest that immunogenicity ultimately depends on the number of detectable TAAs at the time of impaired immune recognition, suggesting that TAA-depleted tumors share in common the tendency for their evasion strategies to incur less antigenic penalties. Our results would predict the possibility to alter the tumor microenvironment to increase the immunogenicity of immune-cold tumors by making evasion more costly, in a manner reminiscent of mutational meltdown [10]. We remark that these dynamics are worth considering in the case of adoptive T cell-based immunotherapies, which have a large potential for exerting substantial co-evolutionary pressure on a developing malignancy [11].

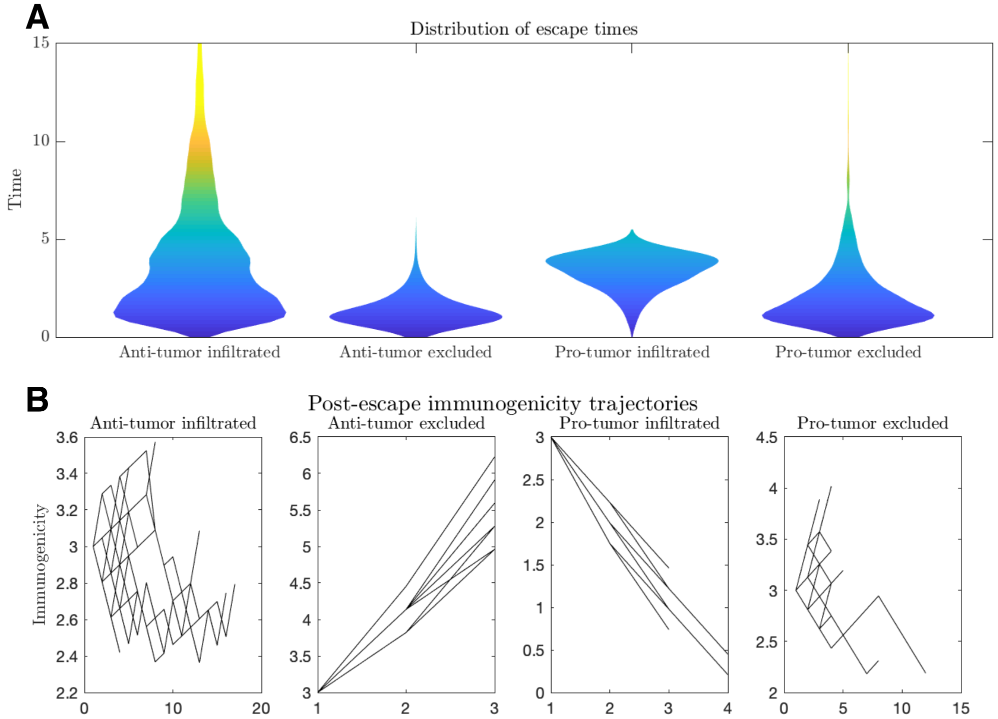


Figure S14: Escape dynamics. (A) Violin plots of the distribution of escape times and (B) Immunogenicity (number of relevant TAAs) as a function of time for a variety of tumor microenvironmental conditions (Anti-tumor infiltrated: $q = q^* + 0.1$, $\beta = 0.529$; Anti-tumor excluded: $q = q^* - 0.1$, $\beta = 0.505$; Pro-tumor infiltrated: $q = q^* + 0.1$, $\beta = -0.529$; Pro-tumor excluded: $q = q^* - 0.1$, $\beta = -0.505$. In all cases, $\beta$ chosen to give $|s^*| = 3$. Simulations were run until $n = 10^6$ escape events occurred for each case. Plot B trajectories were obtained by plotting the immunogenicity trajectories for 50 post-escape events at each time point $n$ for which $n \geq 2$).

### S3.6 Survival benefit of active evasion

From the above analysis, immunogenicity dynamics of an active Evader are closest to those of a mean-reverting passive Evader under the pro-tumor infiltrated case. Given this, we study the dynamics under active and passive evasion as well as the distribution of escape times and probability of escape (Fig. 2). For a reasonable comparison, we fix $q$ and $s^*$ for each case, and the passive evasion rate $p$ is chosen to match the stationary mean optimal evasion rate $\pi^*$. Our simulations result in escape occurring 1.6 times more frequently under active evasion. Moreover, active evasion exhibits a broader distribution of elimination and escape times (Mean Passive Escape = 6.0, Var Passive Escape = 25.0, Mean Passive Elimination = 6.1, Var Passive Elimination = 30.1 ; Mean Active Escape = 7.2 Var Active Escape = 35.8, Mean Active Elimination = 6.7, Var Active Elimination = 38.0). Our results demonstrates that active evasion allows an Evader to adapt to the observed recognition and, despite continual penalty, allows an Evader to 'out-wait' a Recognizer in order to undergo escape.

### S3.7 Exogenous recognition and conclusions

One powerful advantage of this approach is that the theoretical predictions are not limited by the underlying distribution of $r_n$ driving the process. In fact, the optimal policies and value function can handle any temporally varying recognition landscape, $\{r_n\}_n$, so long as $0 \leq r_n \leq s_n$. We consider the effects of step, cyclical, increasing, and decreasing recognition landscapes on the relative evasion probability for populations adopting either a passive or active strategy (Fig. 3).

In addition to arbitrary recognition landscapes, our dynamic programming approach may be applied to understand the effects of immunotherapeutic intervention, whereby immune escape can be modeled as a range of possible behavior on the spectrum of passive evasion to the most aggressive (active) evasion. For example, the active evasion dynamics assuming an anti-tumor infiltrated case are similar to those of passive evasion. In both cases, the process escapes with immunogenicity values that fluctuate around a stationary $s^*$. We can recover the recover the relationship between $s^*$ and mutation rate $\nu(n) = \Delta\lambda/\Delta n$ via Eqs. S25,S39 for the passive case and Eqs. S85,S90 for the active case. In both cases, the result is similar:

$$s^* = \nu/2q. \tag{S102}$$

demonstrating that immunogenicity, and thus the success likelihood of immunotherapeutic intervention, varies directly with mutation rate and inversely with recognition rate. This theory predicts that escape to a cold tumor is more likely when $s^*$ is close to 0 and is akin to complete evasion as modeled in [12], contrasting with temporary evasion that may be recognized subsequently [13]. All else equal, higher mutational rates can lead to higher predicted efficacy via higher $s^*$, but this is not the only way, as concomitantly high rates of recognition can drive $s^*$ down, thereby reducing predicted efficacy." In Eq. S102 above, it is clear that a better immunotherapy prognosis occurs when the mutational rate is higher and the recognition rate is also low, since $s^*$ is predicted large in this case. Fig. S15 summarizes the behavior of an adaptive Evader subject to a temporally varying recognition pressure.

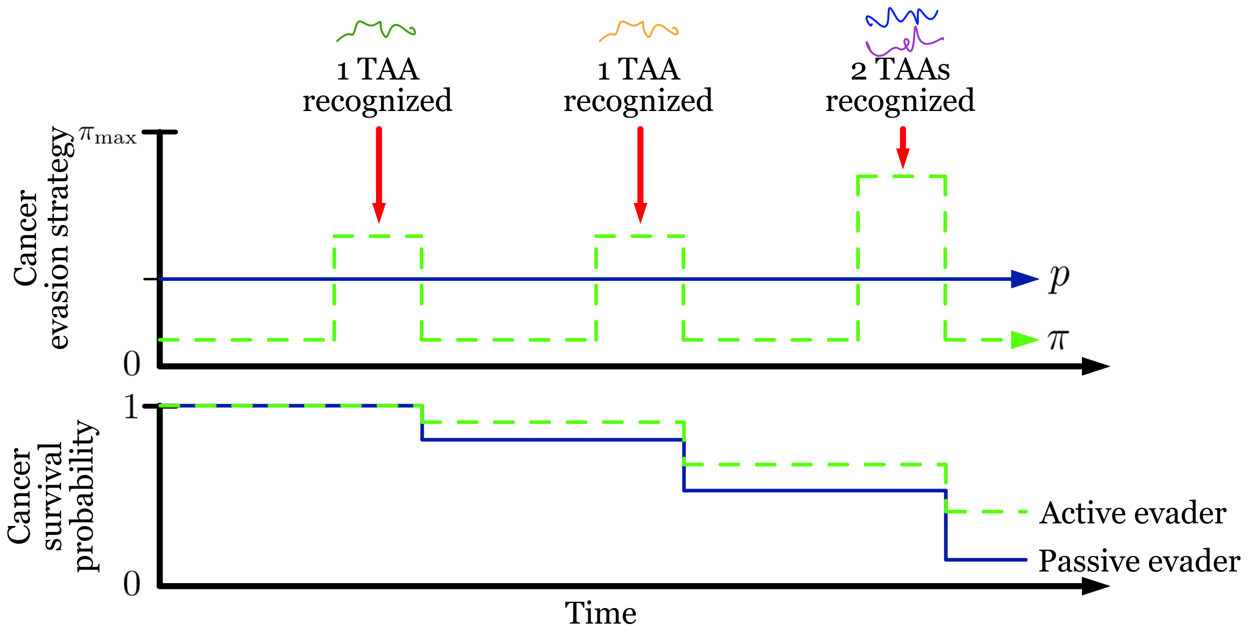


Figure S15: Active evasion summary. Summary of optimized adaptive vs passive evasion on temporally varying recognition and their effect on overall cancer survival.)
